# Phase separation by the Sterile Alpha Motif of Polyhomeotic compartmentalizes Polycomb Group proteins and enhances their activity

**DOI:** 10.1101/2020.08.20.259994

**Authors:** Elias Seif, Jin Joo Kang, Charles Sasseville, Olga Senkovitch, Alexander Kaltashov, Elodie L. Boulier, Ibani Kapur, Chongwoo A. Kim, Nicole J. Francis

## Abstract

Polycomb Group (PcG) proteins organize chromatin at multiple scales to regulate gene expression. A conserved Sterile Alpha Motif (SAM) in the <u>P</u>olycomb <u>R</u>epressive <u>C</u>omplex 1 (PRC1) subunit Polyhomeotic (Ph) is important for chromatin compaction and large-scale chromatin organization. Like many SAMs, Ph SAM forms helical head to tail polymers, and SAM-SAM interactions between chromatin-bound Ph/PRC1 are believed to compact chromatin and mediate long-range interactions. To understand mechanistically how this occurs, we analyzed the effects of Ph SAM on chromatin *in vitro*. We find that incubation of chromatin or DNA with a truncated Ph protein containing the SAM results in formation of concentrated, phase-separated condensates. Condensate formation depends on Ph SAM, and is enhanced by but not strictly dependent on, its polymerization activity. Ph SAM-dependent condensates can recruit PRC1 from extracts and enhance PRC1 ubiquitin ligase activity towards histone H2A. Overexpression of Ph with an intact SAM increases ubiquitylated H2A in cells. Thus, phase separation is an activity of the SAM, which, in the context of Ph, can mediate large-scale compaction of chromatin into biochemical compartments that facilitate histone modification.

## Introduction

Polycomb Group (PcG) proteins repress gene expression by modifying chromatin at multiple scales, ranging from post-translational modification of histone proteins to organization of megabase scale chromatin domains ^1-6^. Two main PcG complexes, PRC1 and <u>P</u>olycomb <u>R</u>epressive <u>C</u>omplex 2 (PRC2), are central to PcG function and conserved across evolution ^1-4^. Both complexes can carry out post-translational modification of histones (methylation of histone H3 on lysine 27 (H3K27me) for PRC2 and ubiquitylation of lysine 118/119 of histone H2A (H2A-Ub) for PRC1. PRC1, and to a lesser extent, PRC2, are also implicated directly in long rage organization of chromatin and clustering of PcG proteins into foci in cells. Two classes of PRC1 complexes have been defined, canonical (cPRC1), and non-canonical (ncPRC1). Both types of complexes contain two ring finger proteins required for E3 ubiquitin ligase activity towards H2A (Psc and dRING in *Drosophila*, Pcgf and Ring1A or B in mammals) ^3,4^. cPRC1 additionally contains a Cbx protein (Pc in *Drosophila*), and a PHC (Ph in *Drosophila*). In ncPRC1, RYBP replaces the Cbx protein, PHCs are absent, other accessory proteins are variably present, depending on the Pcgf subunit ^4^. At least in mouse embryonic stem cells, ncPRC1 is responsible for the bulk of ubiquitylated H2A ^7,8^. This suggests histone modification and chromatin organization may be partitioned between nc and cPRC1s, although both types of complexes share many genomic targets ^7-10^. All cPRC1 subunits can interact with DNA and/or chromatin, and both canonical and non-canonical PRC1s can compact chromatin *in vitro* ^9,11^, but Polyhomeotic (Ph), and thus canonical PRC1, is the most implicated in large-scale chromatin organization ^3,12-17^.

Ph is a core subunit of canonical PRC1, and its most notable feature is the presence of a conserved Sterile Alpha Motif (SAM) in its C-terminus that can assemble into helical polymers ^18^. SAMs are present in many different types of proteins and in many cases can mediate protein polymerization ^19^. The SAM of Ph is required for Ph function in *Drosophila*, and its full polymerization activity is important for gene repression ^20,21^. PRC1 forms visible foci both in *Drosophila* and in mammalian cells^13,22^, and, in *Drosophila* cells, a much larger number of diffraction-limited clusters ^14^. Disrupting the Ph SAM impairs formation of PcG protein clusters and reduces long-range contacts among PcG bound loci, suggesting the two processes are related ^13,14^. Despite the wealth of *in vivo* data supporting the critical function of Ph SAM in large-scale organization of PcG proteins and chromatin, and in gene regulation, the biochemical mechanisms by which Ph SAM links protein and chromatin organization are not known.

In recent years, an important role for liquid-liquid phase separation (LLPS) in organizing macromolecules in cells has been defined ^23-26^. This mechanism is increasingly accepted as being important in formation of protein-RNA membraneless organelles^26,27^, and has more recently been implicated in chromatin compartmentalization and genome organization ^28-33^, transcription activation ^34-37^, DNA repair ^38,39^, and PcG protein organization ^40-42^. LLPS by nuclear/chromatin associated proteins may concentrate proteins and RNAs, enhance or inhibit reactions, exclude other factors, and even physically move genomic regions ^23,25,43^. Nucleated phase separation at super-enhancers mediated by disordered regions in transcription factors and co-activators is believed to be important for driving cycles of active transcription ^34,35^. Phase separation is also implicated in the formation of heterochromatin and its function as a distinct chromatin environment ^30,32^, although the precise role of LLPS is debated ^44^. The mammalian PcG protein Cbx2 (part of certain cPRC1s) has also been shown to undergo LLPS *in vitro* with chromatin, and to form foci in mammalian cells, suggesting a link between LLPS and PcG function ^40,42^.

Here, we consider the hypothesis that Ph SAM can organize chromatin though phase separation by analyzing Ph-chromatin interactions *in vitro*.

## Results

### A truncated version of Ph, “Mini-Ph” forms phase separated condensates with DNA or chromatin

In *Drosophila melanogaster*, the *Ph* gene is present as a tandem duplication in the genome; the two genes (*Ph-p* and *Ph-d*) encode highly related proteins with largely redundant function ^45^. *Drosophila* Ph is a large protein (1589 amino acids for Ph-p), the majority of which is disordered (Fig. 1A), and which is difficult to work with *in vitro*. To focus on the function of the domains conserved in Ph orthologues, particularly the SAM, and to facilitate biochemical analysis, we used a truncated version of *Drosophila* Ph-p, termed “Mini-Ph” ^5^. Mini-Ph (aa1289-1577) contains the three conserved domains—from amino-to carboxy-terminus: the HD1, the FCS Zinc finger that can bind nucleic acids ^46^, and the Ph SAM (Fig. 1A). An unstructured linker connects the FCS to the SAM, and restricts Ph SAM polymerization ^5^. Thus, while Ph SAM alone forms extensive helical polymers *in vitro*, Mini-Ph exists mainly as short polymers of 4-6 units (Fig. 1B-C), even at high concentrations ^5^.

**Figure 1.**
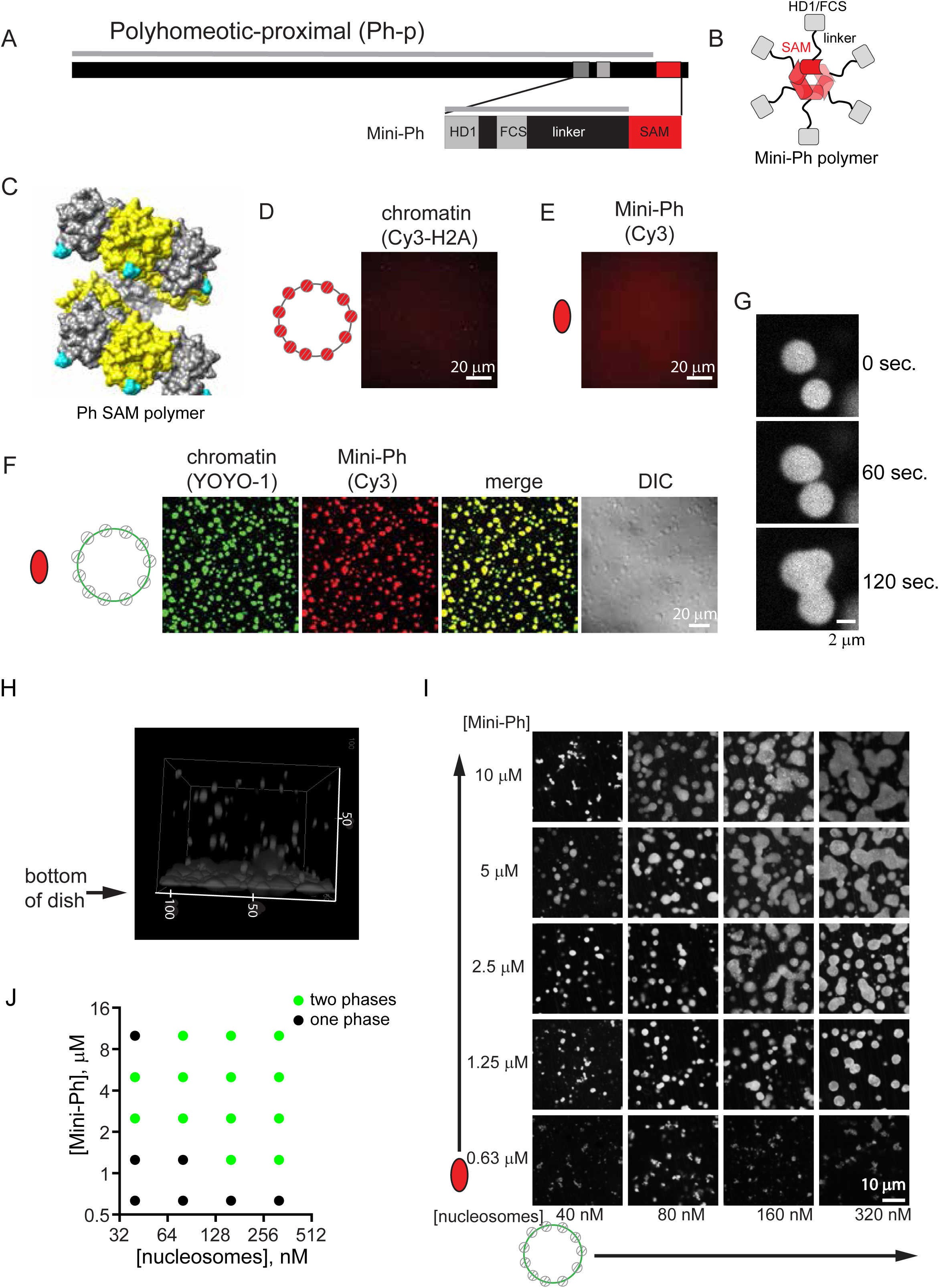
Mini-Ph forms phase separated condensates with chromatin. A. Schematic of Polyhomeotic-proximal and Mini-Ph, which spans aa 1389-1577 and includes the 3 conserved domains and an unstructured linker. Grey line indicates predicted disordered sequence (using PONDR-VSL2)^94^. Note that 91.9% of the sequence is predicted to be disordered (disregarding segments less than 30 amino acids), with only the SAM predicted to be ordered. B. Schematic depicting the oligomeric state of Mini-Ph, which forms limited polymers of 4-6 units (6 are shown) ^5^. C. Structure of nine units of the Ph SAM polymer demonstrating its helical architecture. The N-terminus, from which the linker extends, is shown in cyan. D, E. Neither chromatin (D), nor Mini-Ph (E) form condensates in buffer. F. Mini-Ph forms phase separated condensates with chromatin. G. Time-lapse of droplet fusion of Mini-Ph-chromatin condensates, visualized with Alexa 647-labelled Mini-Ph. H. 3D-reconstruction of confocal stack of images demonstrating that Mini-Ph-chromatin condensates form a fused layer on the bottom of the imaging plate. Scale is in microns. I. Representative images from a matrix of Mini-Ph and chromatin showing the relationship between protein and chromatin concentration and condensate formation. [nucleosomes] assumes 8 fmol nucleosomes per 1 ng DNA. J. Graph depicting conditions where one-phase and two-phases were scored in two experiments like the one shown in I. See also Supplementary Figures 1-3 and Supplementary Movies 1-3.

We expressed Mini-Ph in *E. coli*, purified it (Supplementary Fig. 1A), and tested whether it can form phase-separated condensates, alone or with chromatin. Chromatin was prepared on a circular plasmid containing 40 copies of the *Lytechinus* 5S rDNA nucleosome positioning sequence ^47^ using histones fluorescently labelled with Cy3 on histone H2A (Fig. S1C-E). Neither Mini-Ph alone, nor chromatin alone formed condensates in buffer (Fig. 1D, E). When Mini-Ph is mixed with chromatin, or plasmid DNA, large, round, phase bright drops are observed (Fig. 1F; Supplementary Fig. 2A). Drops formed with either DNA or chromatin undergo fusion (Fig. 1G, Supplementary Fig. 2B; Supplementary Movie 1, 2), and settle to the bottom of the imaging plate where they flatten and continue to fuse (Fig. 1H). Under phase separation conditions, Mini-Ph and DNA can be pelleted by centrifugation (Supplementary Fig. 2C, D), consistent with them forming a denser phase. Mini-Ph-DNA solutions also become turbid, as measured by OD_340_ (Supplementary Fig. 2E). To evaluate the relationship between the concentration of chromatin or DNA and Mini-Ph, and phase separation, we titrated both Mini-Ph and DNA or chromatin, and manually scored each point in the resulting matrix as “one phase” or “two phases” (Fig. 1I, J; Supplementary Fig. 2F, G). This produces a limited coarse-grained delineation of the boundary between one- and two-phase regimes. Phase separation is sensitive to the concentration of both components, and the ratio between the two. This is most notable for Mini-Ph-DNA titrations, where we are able to add high concentrations of DNA, which prevent phase separation (Supplementary Fig. 2F, G). From similar titrations of NaCl and Mini-Ph at a fixed DNA concentration, we find that phase separation is observed in NaCl concentrations up to 125 mM (Supplementary Fig. 3). We conclude that Mini-Ph forms phase separated condensates with either DNA or chromatin.

The disordered linker connecting Ph SAM to the FCS domain was previously demonstrated to restrict Ph SAM polymerization, possibly due to its ability to contact Ph SAM *in trans* ^5^. A scrambled linker has the same effect, implicating amino acid composition rather than organization ^5^. The sequence properties of linkers that connect structured domain play a central role in phase separation ^48^, by restricting or promoting interactions between structured domains, and by contributing weak interactions ^49^. We therefore analyzed the sequence properties of the linker (Supplementary Fig. 4), both in *Drosophila* Ph, and in the three human homologues (PHC1-3). The Ph linker is acidic (pI 3.9), but relatively uncharged (fraction charged residues (FCR) =0.15), and does not have strongly segregated charge (Supplementary Fig. 4B, E, Supplementary Table 1). Overall, the Ph linker is expected to be collapsed (Supplementary Fig. 4D).

The linker region is conserved between the two *Drosophila* Ph homologues (Supplementary Fig. 4F), but both the sequence and charge properties of the linker in mammalian PHCs are distinct (Supplementary Fig. 4C, D, E, G; Supplementary Table 1). The human PHC linkers are basic (pI >10), more charged (FCR 0.25-0.34), have more segregated charges, and have a higher fraction of expansion promoting residues (Supplementary Fig. 4C, E). They occupy a distinct position on the Das-Pappu diagram of states (Supplementary Fig. 4D), predicting context dependent collapse or expansion. Previous analysis indicates that the PHC3 linker promotes polymerization of either PHC3 or Ph SAM, and does not interact with the PHC3 SAM in trans ^50^. A synthetic linker designed to be unstructured (Rlink ^5^) promotes polymerization of both Ph and PHC3 SAM, and shares properties with PHC linkers, including a basic pI (Supplementary Table 1). Evolutionary tuning of the linker sequences is likely to affect phase separation properties of PHCs, although this will need to be tested experimentally.

### Chromatin is highly concentrated in Mini-Ph condensates

One potential function of phase separation is to concentrate (compact) chromatin. To measure the concentration of chromatin in Mini-Ph-chromatin condensates, we first prepared calibration curves using the same Cy3-labelled histone octamers (labelled on H2A) that were used to assemble chromatin (Supplementary Fig. 5A). The concentration of nucleosomes in Mini-Ph condensates, starting from a mixture of 150 nM nucleosomes, and 5 µM Mini-Ph, was measured as 22.5 +/- 4.4 µM (Supplementary Fig. 5B). We note that this value is lower than the reported concentration of chromatin in pure chromatin condensates induced by monovalent cations (∼340 µM ^28^). The reported measurements used free dye to prepare the calibration curve. When we imaged calibration curves prepared from free Cy3, although the curves are linear, they predict at least a 60x higher concentration than curves prepared with labelled histone octamers using the same imaging parameters. Because ladders prepared with free Cy3 do not accurately predict known concentrations of Cy3-labelled histone octamers in our hands, we believe the chromatin concentrations measured using the Cy3-labelled histone calibration curve (Supplementary Fig. 5) are correct for Mini-Ph-chromatin condensates.

### Mini-Ph is dynamic in condensates, but chromatin intermixes slowly

A characteristic of liquid condensates is that the components are dynamic. We carried out fluorescent recovery after photobleaching (FRAP) experiments with Mini-Ph-chromatin condensates. A fraction of Mini-Ph is mobile and exchanges in condensates, so that bleached Mini-Ph drops partially recover fluorescence within several minutes (Fig. 2A, B; Supplementary Fig. 6A-D). In contrast, when the histones (labelled with H2A-Cy3) were bleached, less than 15% of the fluorescence is recovered after several minutes (Fig. 2B, Supplementary Fig. 6E, F). We quantified our FRAP data with user selected ROI for the bleach area and background, and fit the data with a double exponential function. Recent work has drawn attention to the complexity of FRAP measurements in phase-separated condensates, and in selecting and applying the appropriate biophysical model to the data ^51^. Because of the complexities cited above, we interpret the FRAP curves qualitatively. Although we have calculated the half-times of the fast and slow populations and mobile fractions from our data (Supplementary Fig. 6A-D), we do not think these numbers can be used to compare with other systems, or with the measured FRAP behavior of Ph *in vivo* ^52^. Nevertheless, they indicate that Mini-Ph and chromatin have very different kinetics in condensates. Similar behavior has been dissected in a model system of lysine or argnine rich peptides and homopolymers of RNA ^53^. In this case, slow kinetics for the RNAs could be explained by RNA-RNA interactions ^53^. It is possible that nucleosome-nucleosome interactions contribute to the slow kinetics of chromatin. However, it must also be emphasized that the chromatin templates used in these experiments are large (11 kb of DNA, ∼55 nucleosomes, ∼13750 kDa). This system may partially mimic chromatin *in vivo*, which also does not freely intermix (discussed in ^54^). Experiments with 12-nucleosome linear arrays (more than 4X smaller than the templates used here) indicate that while chromatin alone can form a liquid like state that shows (slow) recovery in FRAP experiments ^28^, in most conditions, chromatin forms condensates that behave as solids and do not recover in FRAP experiments, similar to chromaitn *in vivo* ^55^.

**Figure 2.**
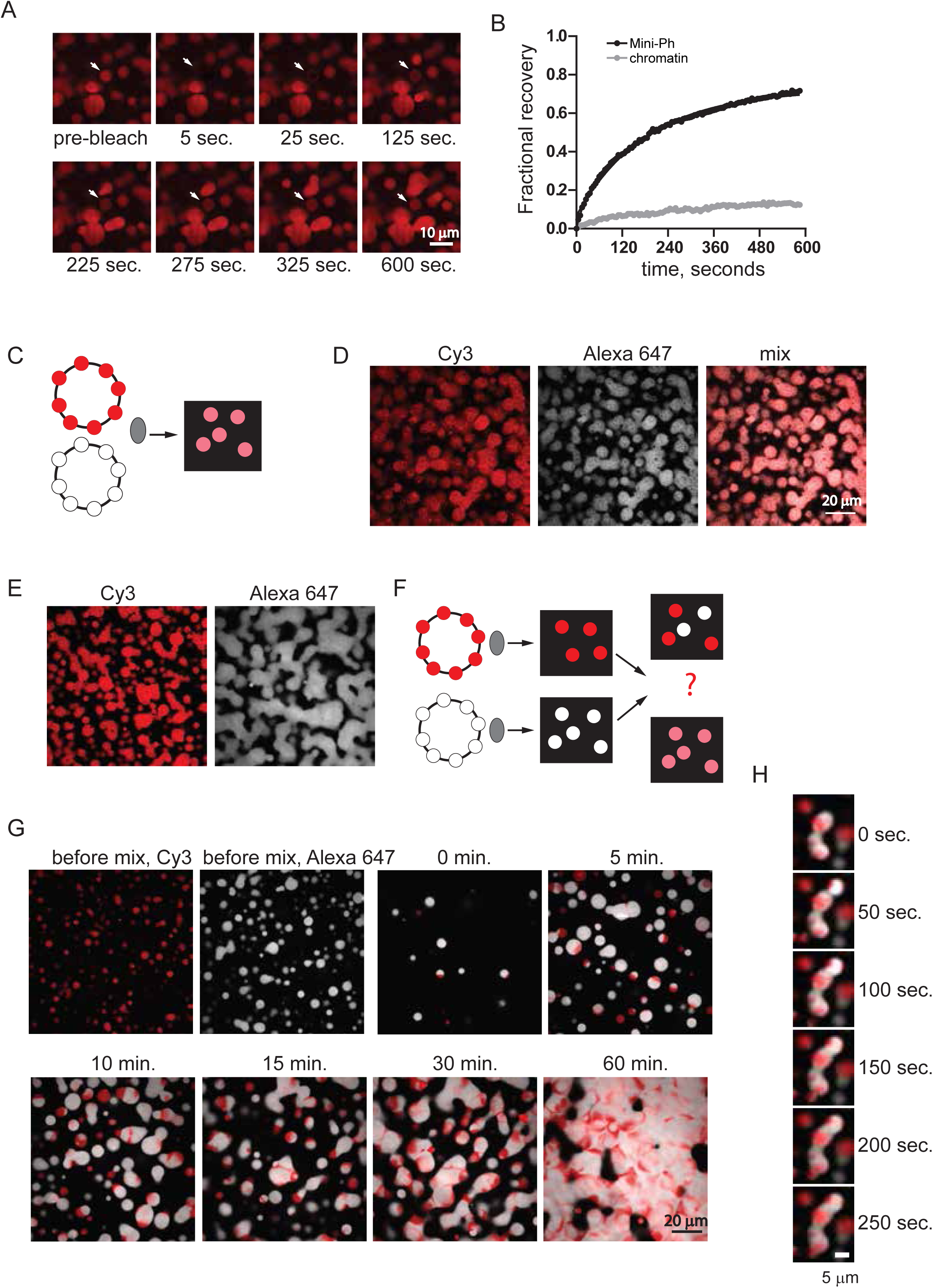
Mini-Ph chromatin condensates intermix slowly *in vitro* although Mini-Ph is dynamic. A. FRAP experiment demonstrating that Mini-Ph-chromatin condensates exchange Mini-Ph. Structure indicated with the arrow was bleached at t=0. B. Representative FRAP traces for Mini-Ph or chromatin (after 28 or 27 min. of condensate formation, respectively). Data were fit with a double exponential function. For Mini-Ph, % fast=25; T1/2_Fast_=30 sec; T1/2_Slow_=199 sec; mobile fraction (plateau)=0.79. For chromatin, % fast=19; T1/2_Fast_=21 sec; T1/2_Slow_=289 sec; mobile fraction =0.14. C. Scheme for mixing chromatin labelled with different colours before adding Mini-Ph. D. Mini-Ph-chromatin condensates formed with an equal mix of Cy3 and Alexa-647-labelled chromatin. E. Mini-Ph-chromatin condensates formed with either Cy3 or Alexa-647-labelled chromatin. F. Scheme for mixing Mini-Ph-chromatin condensates formed with differently labelled chromatins. G. Mini-Ph-chromatin condensates were formed with Cy3- or Alexa 647-labelled chromatin and mixed together. H. Time lapse of fusion of condensates in the mixing experiment. See also Supplementary Fig. 6.

To further understand how chromatin intermixes in condensates, we used two-colour chromatin experiments (Fig. 2C-H). Mini-Ph was incubated separately with chromatin labelled with Cy3 or Alexa 647. Once condensates had formed, the two sets were mixed together, and images collected as the condensates fused (Fig. 2C-H). Although condensates of both colours fused, ultimately forming a fused network at the bottom of the imaging plate (Fig. 2G, H), distinct Cy3 and Alexa-647 regions remained, indicating that the chromatin in pre-formed condensates does not fully intermix when the condensates fuse, at least over the 60 minutes that we monitored (Fig. 2H). This is in clear contrast to control experiments in which the two chromatins are mixed prior to addition of Mini-Ph, where all structures contain a uniform mix of both fluorophores (Fig. 2C, D). These experiments are consistent with the coexistance of different dynamics in Mini-Ph-chromatin condensates. The persistence of unmixed regions could also reflect dynamically arrested phase separation in the pre-formed condensates. We note that in the mixtures shown in Fig. 2C-H, the Alexa-647 labelled chromatin (white in Fig. 2) has a slightly lower nucleosome density than the Cy3 (red) labelled chromatin. The persistent unmixed regions tend to be red regions at the junctions of fused drops. This raises the possibility that nucleosome density affects chromatin dynamics in condensates, due to nucleosome-nucleosome interactions, or nucleosome-Mini-Ph interactions. We conclude that although a fraction of Mini-Ph in Mini-Ph-chromatin condensates is mobile, the chromatin polymers mix slowly and incompletely, a process that could maintain partial compartmentalization of Mini-Ph bound chromatin over short (minutes to hours) time scales.

### Ph SAM, but not its polymerization activity, is required for formation of phase-separated condensates

To test whether Ph SAM is important for condensate formation by Mini-Ph, we prepared Mini-Ph lacking the SAM (Mini-PhΔSAM), or lacking the HD1/FCS domains (Mini-PhΔFCS) (Fig. 3A; Supplementary Fig. 1A). The structure of Ph SAM, including its two polymerization interfaces, termed “End Helix” (EH) and “Mid Loop” (ML) is well characterized ^18^ (Fig. 3B). Mutation of these interfaces blocks SAM polymerization *in vitro* and impairs Ph function *in vivo* ^5,18,20^. We therefore prepared Mini-Ph containing a point mutation that disrupts the EH interface (L1565R) (“Mini-Ph EH”), or a single point mutation that weakens but does not fully disrupt the ML interface (L1547R) (“Mini-Ph ML”) (Supplementary Fig. 1A). Previous AUC experiments with these mutants indicate that Mini-Ph ML forms shorter polymers than Mini-Ph, and Mini-Ph EH at most may form some dimers at high concentrations (see Fig. 3 of ^5^) We first measured the DNA binding activity of each of these proteins using double filter binding with a 150 bp DNA probe (Fig. 3C, D; Supplementary Fig. 7A). Mini-Ph binds DNA with an apparent Kd (Kd_app_) of 37 nM. Partial disruption of polymerization activity with the single ML mutation increases the Kd_app_ to 190 nM. The more severe EH mutation further increases the Kd_app_ to 706 nM, similar to the Kd_app_ of Mini-PhΔSAM (990 nM). DNA binding was not detected with Mini-PhΔFCS by filter binding or EMSA (Supplementary Fig. 7B), indicating that Ph SAM does not bind DNA. Consistent with this conclusion, the Kd_app_ of Mini-PhΔSAM is similar to that for Mini-Ph EH. The much lower Kd_app_ of Mini-Ph presumably reflects cooperative binding by Mini-Ph oligomers. We do not know what the oligomeric state of Mini-Ph is at the concentrations where DNA binding is observed. The Kd_app_ of the SAM-SAM interaction was previously measured as ∼200 nM using an immobilized SAM ^18^, but it is possible that Mini-Ph oligomerization occurs at lower concentrations, which would be consistent with the observed high affinity binding. We conclude that the polymerization activity of Ph SAM increases the affinity of Mini-Ph for DNA.

**Figure 3.**
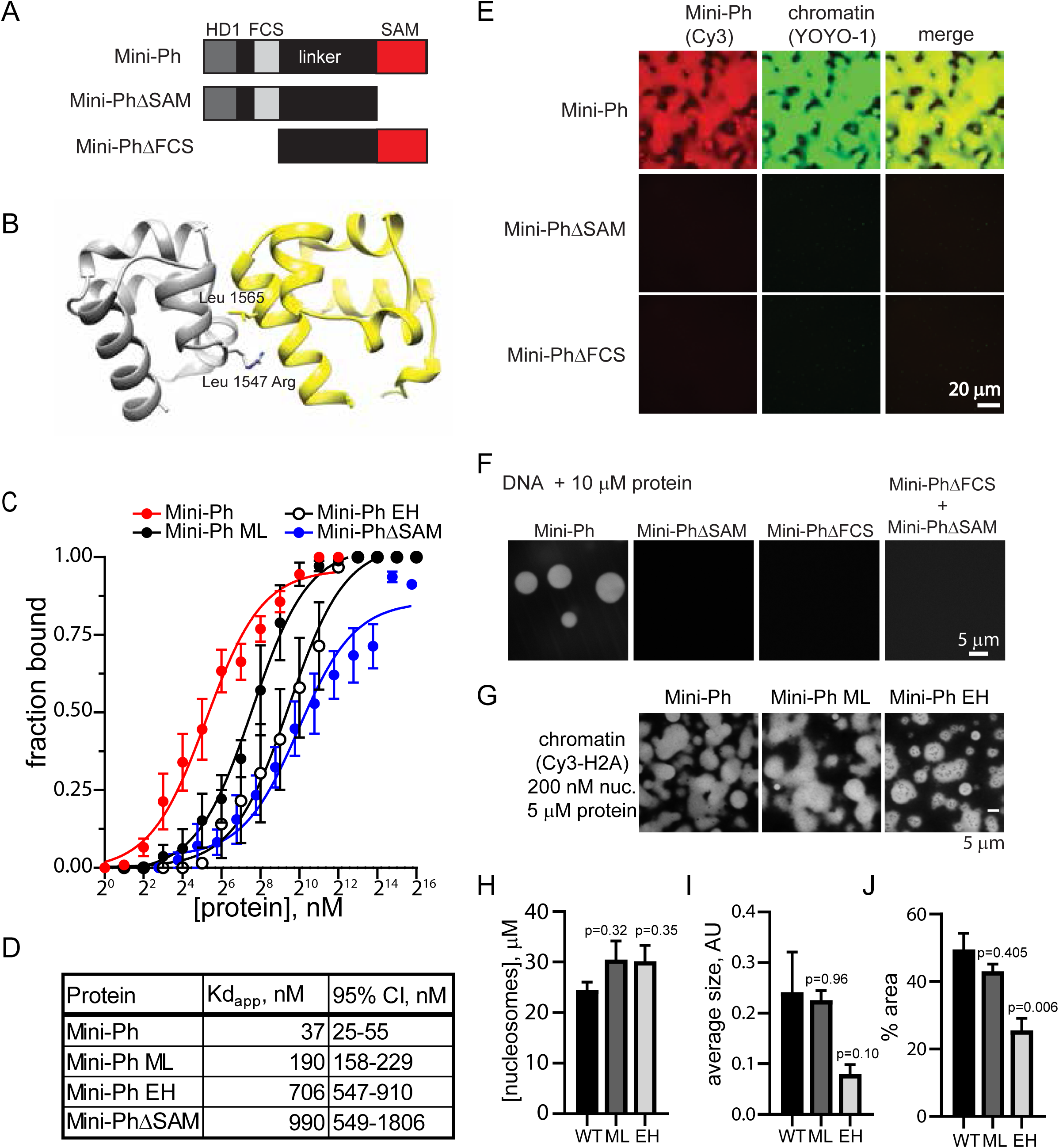
Ph SAM, but not its polymerization activity, is essential for formation of phase-separated condensates *in vitro*. A. Schematic diagram of Mini-Ph truncations. B. Structure of the Ph SAM-SAM interface indicating the position of the ML and EH mutations that impair SAM polymerization. The EH mutation (Leu 1565 Arg) has a stronger effect on polymerization than the single ML mutation (Leu 1547 Arg). The figure was prepared from the structure of the ML mutant, PDB 1D 1KW4. C. Summary of filter binding experiments to measure DNA binding. Points show the mean +/- SEM. D. Kd_app_ for each protein. E. Both the SAM and the FCS/HD1 region are required for formation of phase-separated condensates with chromatin. F. Both the SAM and the FCS/HD1 region are required for formation of phase-separated condensates with DNA or induced by crowding agents. G. Representative images of condensates formed by Mini-Ph or the polymerization mutants (ML and EH) in the presence of chromatin (1 hour incubation). H. The concentration of nucleosomes in condensates formed by wild type (WT) and polymerization mutants (ML and EH) are similar. I, J. EH forms smaller condensates with chromatin than WT Mini-Ph, as determined by measuring the average size of the condensates (I, not significant), or the % area covered by condensates (J). p-values for H-J are for one-way ANOVA comparing each sample to the WT control, with Dunnett’s correction for multiple comparisons. Bars show the mean + SEM of 3 experiments; 9 images were analyzed for each experiment. See also Supplementary Fig. 7.

Neither Mini-PhΔSAM nor Mini-PhΔFCS forms condensates with chromatin or with DNA (Fig. 3E, F). A mixture of the two proteins also does not form condensates with DNA (Fig. 3F). Thus, both the SAM, and the HD1/FCS domains are required for phase separation. We then tested the Mini-Ph polymerization mutants (Mini-Ph ML and Mini-Ph EH). We find that both form phase separated condensates with chromatin or DNA under the same conditions as Mini-Ph (Fig. 3G, Supplementary Figure 8). While the concentration of nucleosomes in condensates is similar (Fig. 3H), condensates formed with Mini-Ph EH are smaller (Fig. 3I, J).

To look more carefully at the effects of the Ph SAM mutations, we titrated Mini-Ph EH or Mini-Ph ML with DNA over a range of NaCl concentrations, and scored each reaction as one-phase or two phases (Supplementary Fig. 8). We find that both mutants are more sensitive to NaCl than Mini-Ph (Supplementary Fig. 3A, B, 8A-C). ATP has been shown to dissolve many protein-RNA condensates, and is hypothesized to have a physiological role in regulating phase separation ^56^. To test whether ATP might also regulate Mini-Ph-chromatin condensates, we formed condensates with Mini-Ph, Mini-Ph ML, or Mini-Ph EH, and challenged them with 2 mM ATP for 15 or 60 min. (Supplementary Fig. 9A). Condensates are smaller after ATP treatment, and Mini-Ph EH is more sensitive than either Mini-Ph or Mini-Ph ML (Supplementary Fig. 9 B-E). Treatment of Mini-Ph condensates with 8 mM ATP completely dissolves them (Supplementary Fig. 10F). We conclude that the Ph SAM, and the HD1/FCS regions are both required for condensate formation, while Ph SAM polymerization activity, which increases DNA binding affinity and changes the oligomeric state of Mini-Ph, enhances condensate formation but is not required for it. Although this result may seem surprising, it is consistent with Mini-Ph existing in a limited oligomeric state prior to condensate formation that cannot be increased further.

Mini-Ph EH and Mini-PhΔSAM have similar DNA binding activities (Fig. 3C), but different abilities to form condensates (Fig. 3D-F). This indicates that the SAM imparts an activity (presumably protein-protein interactions) that is distinct from the effect on DNA binding and polymerization, but essential for condensate formation. The unstructured linker that connects the FCS/HD1 to the Ph SAM (Supplementary Fig. 4A) was previously shown to interact with the SAM (*in trans*) by NMR ^5^. This linker-SAM interaction may allow homotypic interactions between Mini-Ph molecules, even when SAM-SAM interactions are disrupted (as in Mini-Ph EH) and contribute to phase separation. It is also possible that weak SAM-SAM interactions can occur in the EH mutant ^5^ and contribute to phase separation. Ph SAM polymerization may thus indirectly contribute to phase separation by clustering the DNA binding FCS domains (increasing multivalency) and increasing the affinity for DNA/chromatin. Supplementary Fig. 10 summarizes the known and hypothesized interactions that may underlay phase separation by Mini-Ph and DNA or chromatin.

### DNA binding and phase separation modify lysine accessibility in Mini-Ph

To explore how Mini-Ph interactions change on formation of phase separated condensates, and how SAM polymerization affects them, we used a mass spectrometry based protein footprinting method to probe accessible lysines in Mini-Ph (Fig. 4A). We incubated Mini-Ph alone, or with three different amounts of DNA. In the 1X DNA condition (1 Mini-Ph per 10 bp) and 2X DNA conditions, phase separated condensates form, while increasing the DNA amount to 16X prevents their formation (Fig. 4B). To display the data, we generated a heat map of the average accessibility at each lysine under each condition (Fig. 4C). To compare these values, we used student’s t-tests at each lysine position. Accessibility of lysines in the HD1 and FCS domains of Mini-Ph are changed on binding DNA: K1302 and K1340 of the HD1 in Mini-Ph are less accessible in the 2X DNA conditions (condensates present), while K1298 and K1302 are less accessible in the 16X DNA condition (Fig. 4C). As the ratio of DNA to Mini-Ph increases, the accessibility of three lysines in the FCS domain (K1370, K1376, K1380) decreases relative to Mini-Ph alone (Fig. 4C). These decreases in accessibility are consistent with this region being protected by binding to DNA, and indeed, K816 of PHC1 (equivalent to K1380 in Ph) was previously identified as a nucleic acid binding residue in NMR experiments ^46^ (Supplementary Fig. 12A, B). Changes in accessibility could also reflect changes in protein conformation, particularly in the HD1, which is not known to bind DNA. The accessibility of the three lysines in the linker region is low, and does not significantly change with addition of DNA. This is consistent with the linker being in a collapsed state (Supplementary Fig. 4), although the low number of lysines in the linker limits the resolution of the analysis. The accessibility of lysines in the SAM is low both with and without DNA, with no significant changes (Fig. 4C).

**Figure 4.**
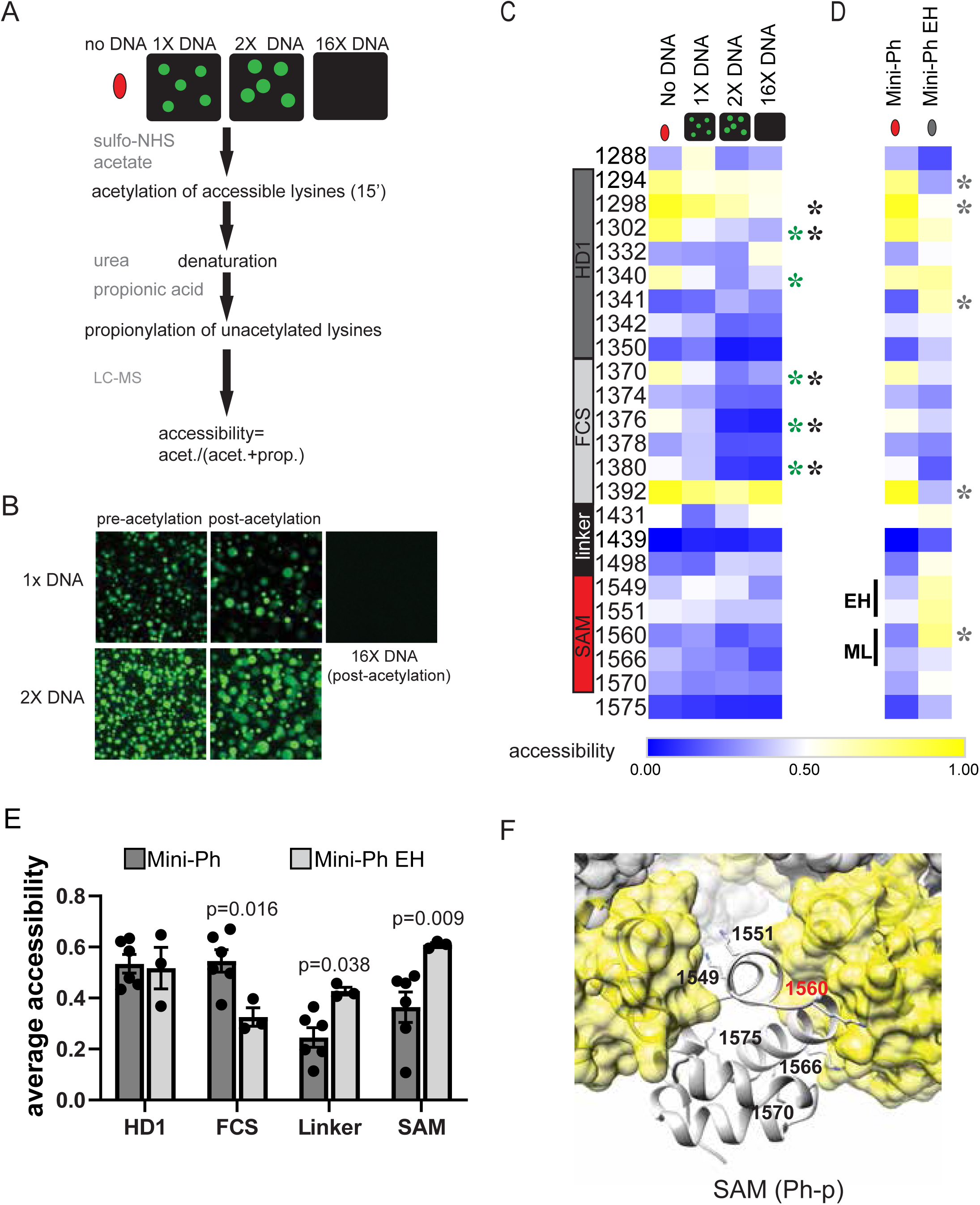
Lysine accessibility in Mini-Ph-DNA condensates. A. Schematic of lysine footprinting assay. Mini-Ph or Mini-Ph EH alone, or in the presence of 1X, 2X, or 16X DNA, is treated with sulfo-NHS acetate to acetylate accessible lysines. The protein is denatured and treated with propionic acid to propionylate unacetylated lysines. Samples are processed for mass spectrometry, and accessibility is quantified as fraction acetylated for each lysine position. B. Mini-Ph-DNA condensates before and after 15 min. acetylation reaction. C. Heat map showing accessibility for Mini-Ph alone or with the indicated DNA amounts (n=3-6). D. Heat map comparing lysine accessibility in Mini-Ph versus Mini-Ph EH (Mini-Ph n=6; Mini-Ph EH n=3). Heat maps are not scaled so that accessibility can be compared across rows and column. Asterisks indicate significant differences between samples with and without DNA by two-tailed student’s t-test with Holm-Sidak correction for multiple comparisons (green=2X DNA vs. no DNA, black =16X DNA vs. no DNA; grey=Mini-Ph versus Mini-Ph EH). E. Average accessibility of lysines in each Mini-Ph region compared between Mini-Ph and Mini-Ph EH. Accessibility of all residues in each region was averaged for each replicate and the averages compared across conditions by student’t t-test with Holm-Sidak correction for multiple comparisons. F. Structure of the Ph-p SAM polymer (PDB 1D 1KW4) with lysine side chains shown and labeled for the central SAM unit. Red highlights residue with significantly changed accessibility in Mini-Ph versus Mini-Ph EH. Structural data are not available for the HD1 residues studied in the footprinting assay. See also Supplementary Fig. 11-13.

To validate global changes in accessibility, we also compared average accessibility of all lysines in each domain under the different conditions (Supplementary Fig. 11C). This confirms the reduction in accessibility of the HD1 and FCS under conditions where condensates form, and no change in the accessibility of the SAM (Supplementary Fig. 11C). These data are consistent with the SAM maintaining its folded structure and pre-existing polymeric state on binding DNA and in condensates. Prolonged incubation in sulfo-NHS-acetate leads to dissolution of condensates (Suplementary Fig. 12 A-C), likely by disrupting binding of Mini-Ph to DNA. Indeed, if Mini-Ph is fully acetylated with sulfo-NHS-acetate, it does not bind DNA, and does not form condensates with DNA (Supplementary Fig. 12 D-F).

We then compared accessibility of lysines in Mini-Ph to that in Mini-Ph EH, which does not form polymers. The pattern of lysine accessibility in Mini-Ph EH is distinct from that of Mini-Ph, and differences are not restricted to the SAM (Fig. 4D). Three lysines in the HD1, one in the FCS, and one in the SAM are significantly altered in Mini-Ph EH versus Mini-Ph. When differences are considered over each domain, they are more striking (Fig. 4E). While the overall accessibility of the HD1 is the same between the two, probably because both increases and decreases in accessibility are observed, the FCS is less accessible in Mini-Ph EH than in Mini-Ph. while the linker and the SAM are more accessible (Fig. 4D-F). The accessibility of the SAM is consistent with the expected monomeric state of Mini-Ph EH and the positions of the lysines in the SAM polymer structure (Fig. 4F). However, thedifferences in the other domains of Mini-Ph EH versus Mini-Ph indicate that SAM polymerization likely affects the whole conformation of Mini-Ph and the interactions available for phase separation. The changes in the HD1 both on condensate formation and between Mini-Ph and Mini-Ph EH also raise the possibility that this domain contributes interactions to phase separation, which will need to be directly tested. Whether Ph SAM would also affect the conformation of Ph in the context of the full length protein, or when it is in PRC1 (an interaction mediated by the HD1) remains to be determined. Finally, we attempted to analyze lysine accessibility in Mini-Ph EH condensates (Supplementary Fig. 13), but the condensates dissolved within 5 min. of adding the acetylation reagent. 5 min. of acetylation results in most of the protein being inaccessible (Supplementary Fig. 13 C, D). After 15 min. of acetylation, accessibility is similar with and without DNA (Supplementary Fig. 13E), consistent with DNA binding being completely disrupted. Comparison of Mini-Ph EH alone after 5 min. or 15 min. of acetylation indicates that the linker and SAM are increasingly acetylated with time (Supplementary Fig. C, F). This is consistent with SAM-SAM and/or linker-SAM interactions (Supplementary Fig. 10), although other explanations are possible.

### Ph SAM Polymerization affects the mobility of Mini-Ph in condensates

The experiments presented above indicate that Ph SAM polymerization increases the DNA binding affinity of Mini-Ph (Fig. 3C), increases the driving forces for phase separation (Fig. 3F-J, Supplementary Fig. 8, 9), and changes the accessibility of Mini-Ph (Fig. 4). To determine whether the polymeric state of Mini-Ph also affects the material properties of condensates, we compared Mini-Ph and Mini-Ph EH mobility in condensates formed with chromatin (Supplementary Fig. 14). In side-by-side experiments, recovery of fluorescence is consistently faster with Mini-Ph EH (Supplementary Fig. 14 A, C, D). We fit FRAP data to a double exponential function. The T1/2 for both slow populations is lower for Mini-Ph EH than for Mini-Ph the % of molecules in the fast fraction is higher for Mini-Ph EH than for Mini-Ph, and the mobile fractions are similar for both (Supplementary Fig. 14 G-J). To analyze chromatin mobility, we analyzed the Cy3 label on H2A in same condensates used to collect FRAP traces for Alexa-647 labelled Mini-Ph or Mini-Ph EH before and after bleaching (Supplementary Fig. 14 B, E, F). Less than 10% of the fluorescence is recovered over the 5 min. experiment for condensates formed with Mini-Ph and Mini-Ph EH (Supplementary Fig. 13B). Thus, consistent with Fig. 2, chromatin and Mini-Ph or Mini-Ph EH have distinct kinetics in condensates. The slow kinetics of chromatin may be intrinsic to the template since the EH mutation in Mini-Ph does not affect them. We conclude that assembly of Mini-Ph into polymers not only increases the driving force for phase separation, but influences the material properties of the condensates that are formed.

### Mini-Ph-chromatin condensates recruit PRC1 from nuclear extracts

One function of phase separation is to create biochemical compartments that are enriched for specific components, and can stimulate or inhibit biochemical reactions. To determine whether Mini-Ph-chromatin condensates can create unique biochemical compartments, we asked whether condensates can recruit proteins from nuclear extracts (Fig. 5A). We prepared nuclear extracts from *Drosophila* S2R+ cells, and used an anion exchange resin to deplete nucleic acids from the extracts. Even after depletion, the nuclear extracts contain substantial amounts of RNA (Supplementary Fig. 15A). Treatment of extracts with RNAseA resulted in precipitation of most of the protein from the extracts, so that we used extracts containing RNA for our experiments. Chromatin alone forms a few tiny structures in extracts (Fig. 5B, C, reaction 1). Mini-Ph does not form condensates in buffer (e.g. Fig. 1C), but does form small condensates in extracts, likely by binding to RNA, since the condensates stain with YOYO-1 (Fig. 5B, C, reaction 2). When Mini-Ph is incubated with chromatin to form condensates, and then nuclear extracts are added, the condensates are preserved, although they are smaller than condensates in equivalent reactions incubated in buffer (Fig. 5B, C, compare reactions 3 and 4). Although the condensates are smaller, the concentration of chromatin in them is similar to that in condensates incubated in buffer (Fig. 5D). We do not know why the condensates are smaller after incubation in nuclear extracts. Post-translational modifications can influence phase separation ^57^, but the small molecule substrates needed for enzymes that mediate them should be depleted in our desalted extracts. The presence of nucleic acids, in the extracts could disrupt condensates, analogous to what is observed at high concentrations of DNA (Supplementary Fig. 2F, G). Alternatively, proteins in the extracts that bind to Mini-Ph and/or chromatin may disrupt interactions required for condensates.

**Figure 5.**
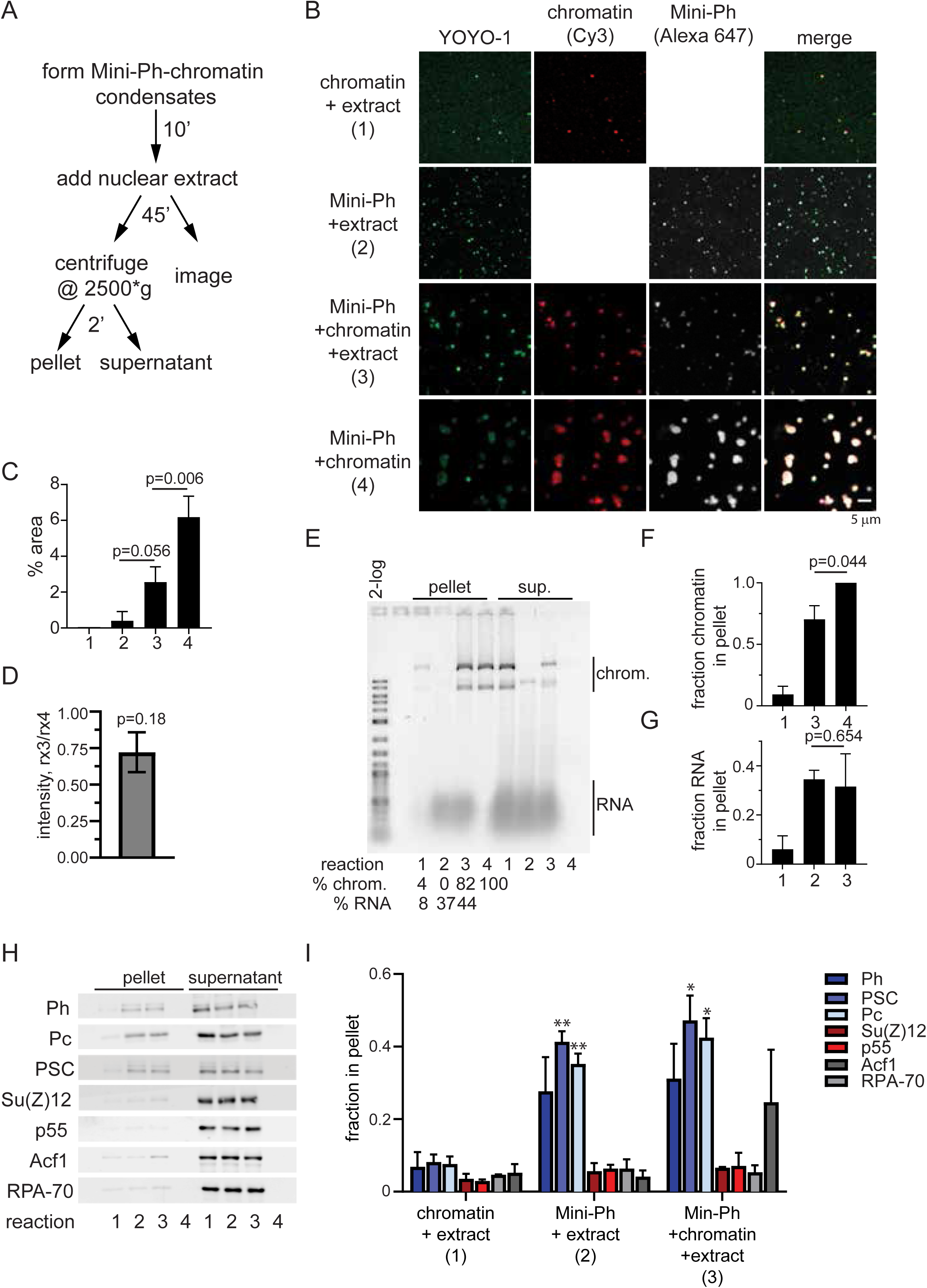
Mini-Ph condensates recruit PRC1 from extracts. A. Scheme for isolating Mini-Ph-chromatin condensates from nuclear extracts. B. Representative images of condensates formed in each of the 4 indicated reactions. C. Quantification of phase separated condensates (% area covered by condensates, 9 images analyzed for each of 3 experiments using YOYO-1 staining). p-values are for one-way ANOVA with Tukey’s correction for multiple comparisons. D. Ratio of average intensity in condensates formed by Mini-Ph + chromatin + nuclear extracts (reaction 3) versus Mini-Ph + chromatin (reaction 4) for 3 experiments. p-value is for one sample t-test comparing the ratio to the expected value of 1. E. SYBR Gold stained gel of nucleic acid content of pelleted reactions. Reactions 1-4 are as indicated in panel B for C-I. Summary of three experiments quantifying the fraction of the chromatin (F) and RNA (G) in the pellet. p-values are for paired t-test between reactions 3 and 4. H. Representative Western blots of one experiment analyzing the content of pelleted condensates. Equal amounts of pellet and supernatants were loaded. I. Summary of three experiments analyzing the content of condensates formed in extracts. Su(Z)12 was only analyzed in two experiments. One-way ANOVA was used to compare all three samples for each antibody with Tukey’s multiple comparisons test. p<0.05 *; p<0.01**. All bar graphs show mean, and error bars are SEM. See also Supplementary Fig. 15.

We used low speed centrifugation (2 min. @ 2500*g) to isolate condensates and analyzed their nucleic acid content on agarose gels. When Mini-Ph is incubated with extract in the absence of chromatin, the pelleted condensates contain RNA (Fig. 5E). When Mini-Ph is incubated with chromatin, and extract added subsequently, the isolated condensates contain both chromatin and RNA (Fig. 5E). Since the amount of RNA that is pelleted with Mini-Ph is similar with and without chromatin, we infer that Mini-Ph condensates in extracts can contain *both* RNA and chromatin (Fig. 5E-G). To confirm this, we analyzed co-localization of fluorescently labelled Mini-Ph with chromatin after incubation in buffer, or in nuclear extracts (Supplementary Fig. 15 B-D). Most Mini-Ph-containing structures also contain chromatin. This is consistent with chromatin condensates recruiting RNA from the extracts, rather than formation of a separate class of Mini-Ph-RNA condensates.

To analyze the protein components of Mini-Ph condensates in nuclear extracts, we used Western blotting. PRC1 components are enriched in condensates formed in extracts with or without chromatin, while the PRC2 subunits Su(Z)12 and p55, the single strand DNA binding protein RPA70, and the chromatin remodeling complex subunit ACF1 are not enriched (Fig. 5H, I). Thus, Mini-Ph condensates can concentrate endogenous PRC1 provided by nuclear extracts.

### Mini-Ph-chromatin condensates enhance ubiquitylation of histone H2A

To determine whether the PRC1 recruited to Mini-Ph condensates is active, we tested whether chromatin present in condensates is ubiquitylated on histone H2A. When extracts were supplied with ATP and ubiquitin, very low levels of ubiquitylated H2A (H2A-Ub) were detected. Addition of the E1 ubiquitin activating enzyme and E2 ubiquitin conjugating enzyme along with ATP and ubiquitin resulted in detectable H2A-Ub in extracts (Fig. 6A, B). Formation of Mini-Ph-chromatin condensates prior to incubation in extracts increased H2A-Ub by about two-fold. This suggests the PRC1 recruited to condensates is functional, and that Mini-Ph-chromatin condensates enhance the ubiquitylation reaction (Fig. 6B, C).

**Figure 6.**
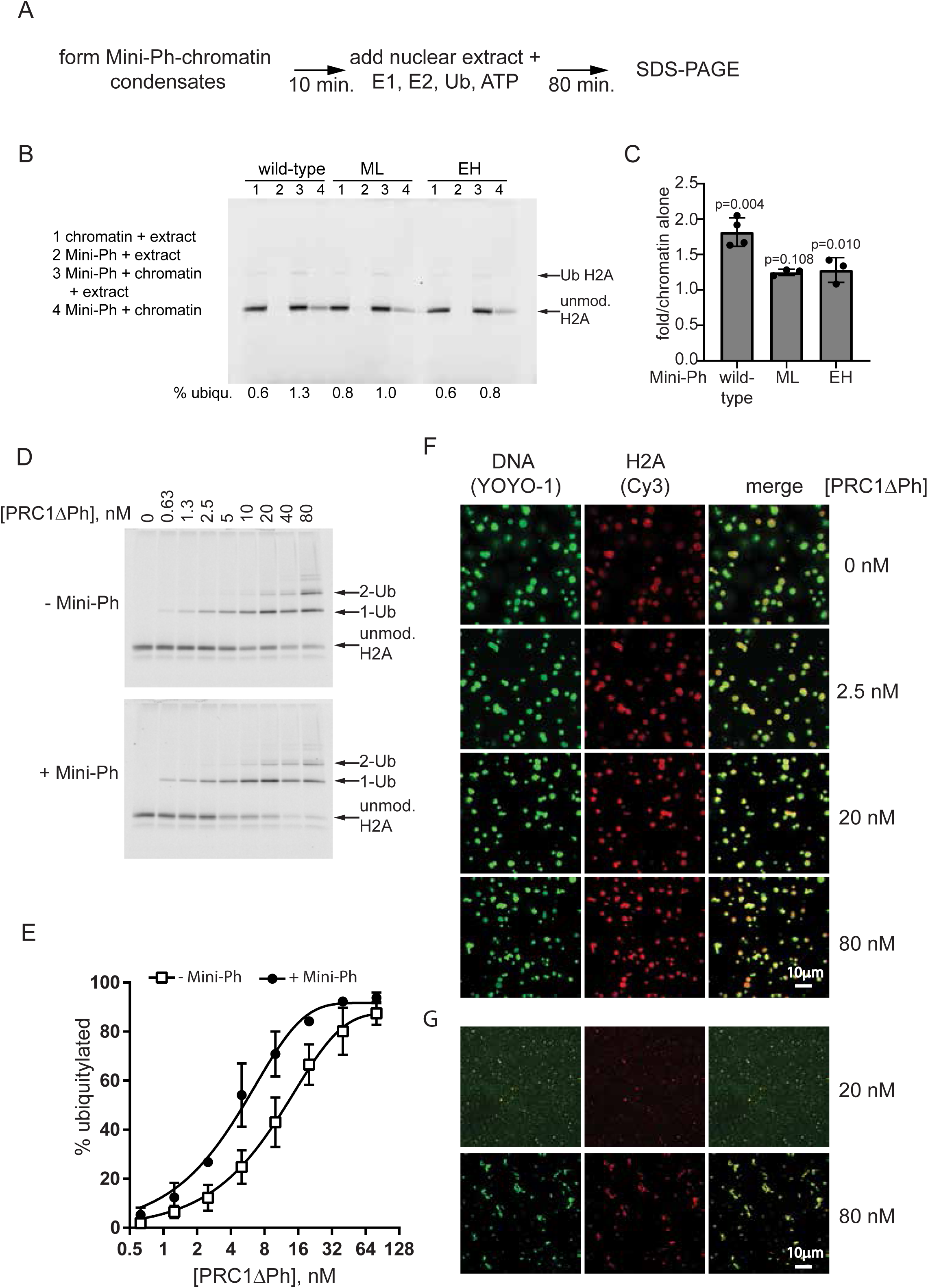
Mini-Ph condensates facilitate histone ubiquitylation by PRC1. A. Scheme for carrying out ubiquitylation assays in nuclear extracts. B. Representative SDS-PAGE of Cy3-labelled H2A showing ubiquitylation of chromatin in nuclear extracts in the presence or absence of Mini-Ph (wild-type), Mini-Ph ML or Mini-Ph EH. Note that the condensates formed in buffer (reaction 4) were poorly recovered in this experiment. C. Quantification of three ubiquitylation assays. Bars are mean + SEM. p-values are for one-sample t-test comparing values to expected value of 1. D. Representative SDS-PAGE of Cy3-labelled H2A showing ubiquitylation reaction with PRC1ΔPh and chromatin in the presence or absence of Mini-Ph. E. Quantification of ubiquitylation reactions; n=3. Points are the mean +/- SEM and data were fit with an exponential function. F. Microscopy of Mini-Ph-chromatin condensates at the end of fully reconstituted *in vitro* ubiquitylation reactions. G. Fibrous condensates are formed by high concentrations of PRC1ΔPh in the absence of Mini-Ph; at lower concentrations, no structures are visible. See also Supplementary Fig. 16, 17.

To determine if the Ph SAM polymerization state can influence condensate formation in the more physiological environment of nuclear extracts, we prepared condensates with Mini-Ph ML, or Mini-Ph EH, and added nuclear extracts to them. Mini-Ph ML condensates behave similar to those formed with Mini-Ph in extracts (Supplementary Fig. 16). In contrast, incubation of Mini-Ph EH condensates in extracts transforms them into diffuse structures that occupy a larger area but have a reduced chromatin concentration relative to condensates incubated in buffer (Supplementary Fig. 16). We tested histone ubiquitylation in extracts in the presence of Mini-Ph ML or Mini-Ph EH, and find that neither mutant stimulates histone ubiquitylation (Fig. 6B, C). We do not know if this is because the condensates formed by the polymerization mutants have different properties (e.g. Supplementary Fig. 14), or because they recruit less PRC1, as would be expected if SAM-SAM interactions (between Mini-Ph and Ph in PRC1) are directly involved in recruiting PRC1 to chromatin.

The observation that Mini-Ph condensates increase histone ubiquitylation might reflect the increased concentration of PRC1 in condensates (Fig. 5H, I). It is not necessarily predicted, however, that the environment of condensates, in which chromatin is compacted, would enhance enzyme activity. Thus to determine whether Mini-Ph-chromatin condensates enhance PRC1 activity under optimal conditions, we reconstituted the ubiquitylation reaction *in vitro*, using chromatin alone or Mini-Ph-chromatin condensates as the substrate (Supplementary Fig. 17). We used PRC1ΔPh for these experiments (Supplementary Fig. 1B), which can interact with Mini-Ph via the HD1 domain (but unlike PRC1 found in extracts, not via SAM-SAM interactions), and is fully active as an E3 ligase. PRC1ΔPh catalyzes formation of H2A-Ub on chromatin in a dose dependent manner (Fig. 6D; Supplementary Fig. 17C, D). When Mini-Ph-chromatin condensates are used as the substrate, the activity of PRC1ΔPh is increased by about two-fold over the entire titration, indicating that condensates stimulate PRC1ΔPh activity (Fig. 6D, E). We also analyzed condensates at the end of the reactions to confirm that they persist under reaction conditions (Fig. 6F, G). Because a high fraction of the histones is ubiquitylated in these experiments (Fig. 6D, E), these results indicate that H2A-Ub does not disrupt condensates.

### Ph SAM affects ubiquitylation of H2A *in vivo*

To test whether the activity of Ph SAM is important for histone ubiquitylation *in vivo*, we used *Drosophila* S2 cell lines that express Ph or Ph with the strong ML mutation (L1547R/H1552R), which disrupts Ph SAM polymerization as effectively as the EH mutant used in our *in vitro* studies, under control of an inducible promoter ^14^. We isolated histones from control S2 cells and cells induced to overexpress Ph or Ph-ML, and measured levels of H2A-Ub (Fig. 7A-D). Cells overexpressing Ph have an approximately two-fold increase in overall H2A-Ub relative to control cells (Fig. 7B). Cells overexpressing Ph-ML have increased H2A-Ub in some experiments, but this difference was not significant, even though Ph-ML is expressed at higher levels than Ph (Fig. 7A, C).

**Figure 7.**
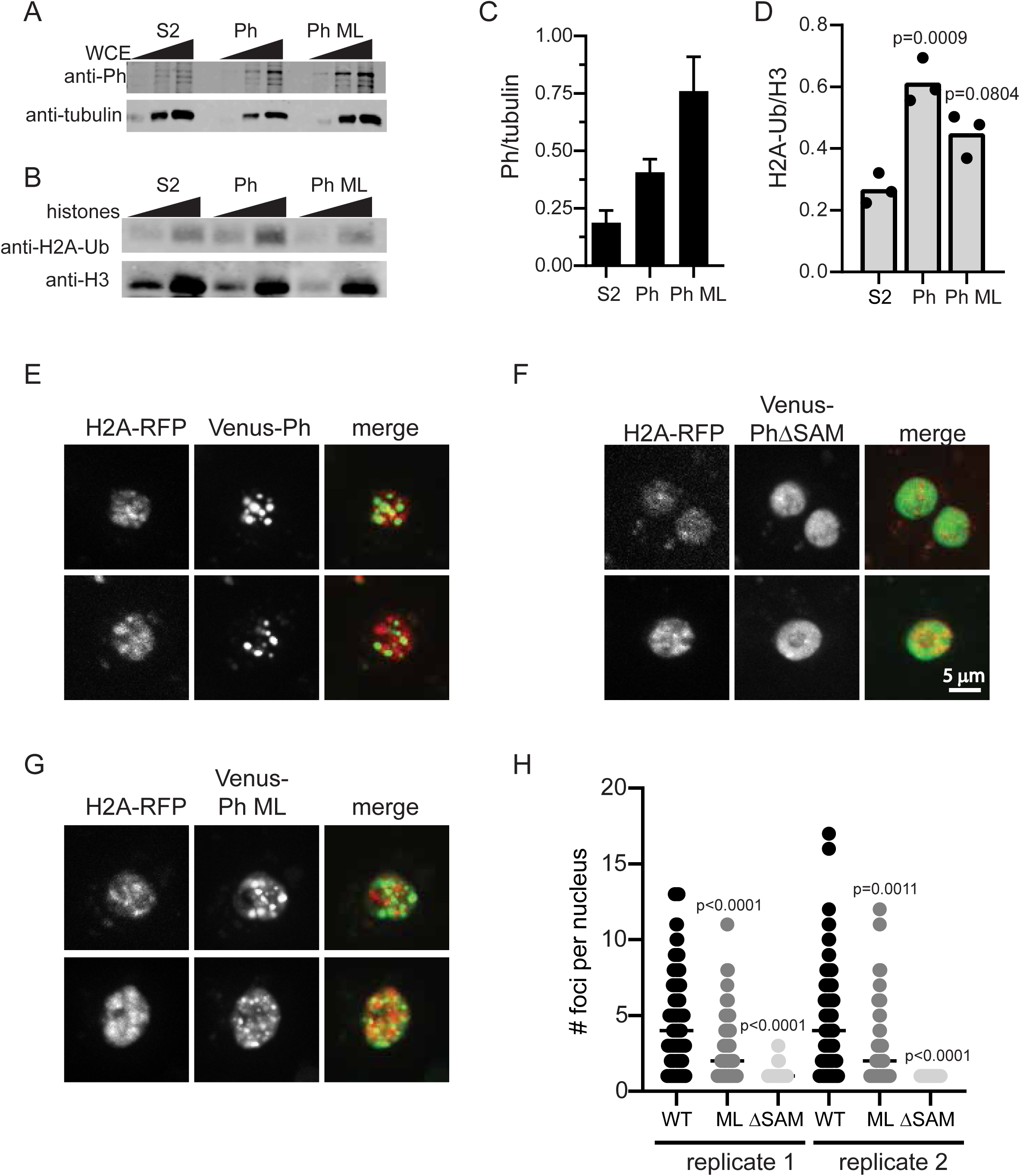
Ph with an intact SAM increases H2A-ub *in vivo*. A. Representative western blot of Ph levels in induced cell lines. Note that Ph ML is the strong double mutant (L1547R/H1552R), which was previously shown to disrupt Ph clustering in cells ^14^. B. Representative Western blot of histone H2A-Ub levels in induced cell lines. Blots were re-probed with ant-H3 to normalize loading. 125 and 250 ng of acid extracted histones were loaded for each sample. C, D. Quantification of Ph (C) and H2A-Ub (D) for 3 experiments. p-values are for one-way ANOVA comparing Ph-WT and Ph-ML cells to control (S2) for the 250 ng point. E-G. Representative images of cells overexpressing YFP-Ph (E), YFP-PhΔSAM (F), or YFP-Ph-ML (G). H. Graph of the number of foci per cell for two independent experiments. Note that only cells with >zero foci were included. p-values are for Kruskal-Wallis tests with Dunn’s multiple comparisons correction. Supplementary Fig. 18.

Because we find that Ph SAM polymerization activity is not strictly required for phase separation *in vitro*, we wondered if Ph-ML might be able to phase separate *in vivo*, particularly when present at high concentrations. Formation of highly concentrated foci in cells is consistent with phase separation, although it can arise through other mechanisms, as has been pointed out ^35^. To test whether Ph-ML can form foci in cells, we transiently transfected *Drosophila* S2 cells with Venus-tagged Ph, Ph-ML, or PhΔSAM under control of the heat shock promoter. After heat shock induction, Venus-Ph forms large, round, bright foci. These foci are mainly (although not exclusively) nuclear, and little Venus signal is observed in the nucleoplasm outside the foci (Fig. 7E). In contrast, Venus-PhΔSAM is uniformly distributed in the nucleus, and does not form foci (Fig. 7F). Venus-Ph-ML forms foci but is also distributed throughout the nucleus (Fig. 7G). Thus, foci formation *in vivo* and phase separation *in vitro* are correlated with each other and with enhanced histone ubiquitylation. We tested Venus-Mini-Ph in *Drosophila* S2 R+ cells, and find that, unlike Venus-Ph, it does not form foci in most cells. In about 7% of the cells, it forms a single focus, which can be quite large (Supplementary Fig. 18 A, B, D); these unusual foci are not observed with Venus-Mini-PhΔSAM (Supplementary Fig. 18C, D). Thus, although Ph SAM is required for foci formation in cells, the other disordered regions of Ph shape its behavior in cells as been observed for other proteins that can undergo LLPS ^57^.

## Discussion

We have identified phase separation as a new activity of the Ph SAM, the domain that is most clearly implicated in large-scale chromatin organization by PcG proteins. Our data are consistent with two possible functions of Ph SAM-dependent phase separation: 1) formation of a compacted yet fluid chromatin state; 2) creating unique biochemical compartments that enhance PRC1-mediated histone modification.

### Phase separation by Ph SAM does not strictly require its polymerization activity

In developing *Drosophila* embryos, Ph lacking the SAM cannot rescue any Ph functions, while Ph with a polymerization interface mutated can partially rescue Ph function, although with defects in transcriptional repression ^20^. *In vitro*, Mini-Ph lacking the SAM does not form phase-separated condensates, while Mini-Ph with the polymerization interface mutated (Mini-Ph EH) does form condensates although they are smaller. In *Drosophila* tissue culture cells, Ph lacking the SAM does not form foci, while polymerization defective Ph (Ph-ML) can form foci when overexpressed (Fig. 7). Thus, foci formation *in vivo*, and phase separation *in vitro* are correlated with full Ph function, and the LLPS activity of Ph SAM may be the critical function of the SAM that remains even when polymerization is disrupted.

Previous work, implicates Ph polymerization in both transcription repression and chromatin organization ^5,13,14,20,21^. <u>S</u>tochastic <u>O</u>ptical <u>R</u>econstruction <u>M</u>icrocsopy (STORM) analysis of PcG proteins in normal *Drosophila* tissue culture cells or those that mildly overexpress Ph or Ph-ML showed that normal *Drosophila* tissue culture cells contain hundreds of nanoscale PcG clusters, although only a few large PcG bodies are visible by conventional microscopy ^14^. Mild overexpression of Ph increased the number but not the size of clusters, and increased long-range contacts, while overexpression of the strong Ph-ML mutant disrupted clusters and reduced long-range contacts. This work and work in mammalian cells ^13^ directly implicates Ph SAM polymerization in the nanoscale organization of PcG proteins and large-scale organization of chromatin.

Although Ph SAM alone can form open-ended polymers, the extent to which long SAM polymers occur in the context of the full protein is unclear. *In vitro*, the oligomeric state of Mini-Ph is limited to four to six units ^5^; this can be explained by the action of the unstructured linker that separates Ph SAM from the FCS in conjunction with the helical configuration of SAM polymers ^18^. Steric considerations suggest Ph SAM polymerization may be even further restricted in the context of PRC1. Thus, the contribution of polymerization to LLPS may be much subtler than would occur with an actual open-ended Ph SAM polymer. The linker connecting the SAM to the FCS is not conserved in Ph homologues (Supplementary Fig. 4). The linker of PHC3, unlike the *Drosophila* Ph linker ^5^, does not bind the PHC3 SAM in trans ^50^, and allows much more extensive SAM polymerization than that of Ph ^50^. It is therefore possible that the linker has been tuned across evolution to control polymerization and its interplay with phase separation. This is consistent with modeling based analysis indicating that the properties of linkers connecting interacting domains tune phase separation properties ^48^. Two other PcG proteins, SCM and Sfmbt, also have SAMs, and the three SAMS have been shown to co-assemble ^58^; joining of SAM-mediated polymers of these three proteins could allow formation of large and diverse polymers. Evaluating the phase separation activity of these other PcG SAMs, alone or in combination, and of Ph homologues, will be an important future goal.

The phase separation activity of Ph SAM is also likely subject to negative regulation. A disordered, serine/threonine rich sequence adjacent to the HD1 undergoes O-linked glycosylation mediated by the PcG protein Sxc ^20,59^. This region, and Sxc, are both important for Ph function in regulation of some genes ^20,59^. In the absence of glycosylation, Ph undergoes SAM-dependent “non-productive aggregation”, which is not alleviated by mutating the Ph SAM polymerization interfaces ^20^. It is possible that “non-productive aggregation” in fact reflects SAM-dependent phase separation (or maturation of phase-separated protein into stable, insoluble aggregates) ^23^. The glycosylated sequence is not part of Mini-Ph. Mini-Ph is produced in *E. coli*, and is not glycosylated, yet Mini-Ph is soluble. It therefore seems likely that the effect of glycosylation, although dependent on Ph SAM, also involves other sequences in Ph. We speculate that the glycosylated region may restrict Ph SAM-mediated phase separation, and preliminary *in vitro* data support this idea (unpublished observation).

A hallmark of LLPS is that it depends on weak, multivalent interactions that allow rapid reorganization and unrestricted stoichiometry. The polymerization activity of Ph SAM may contribute multivalent interactions. However, additional interactions are required, which (at least in Mini-Ph) involve the HD1 and/or the FCS. Based on the comparison between Mini-Ph and Mini-Ph EH, linker-SAM and (possibly) SAM-SAM interactions that do not require an intact polymerization interface likely also contribute (Supplementary Fig. 10). *In vitro*, dynamic SAM polymerization is not likely to directly drive phase separation by Mini-Ph because the Kd for polymerization is so much lower than the saturation concentration at which phase separation occurs. However, in the polymerization mutants, and *in vivo* where the concentration of Ph is lower ^52,60^, dynamic polymerization of Ph SAM could control phase separation. In LLPS of Mini-Ph with chromatin or DNA, the role of the FCS is likely nucleic acid binding; however, the HD1 and/or the FCS may form additional protein-protein interactions (Fig. 4). It is interesting to note that Sfmbt and SCM, the other two SAM containing proteins also contain an FCS, although the distance and additional motifs separating the FCS from the SAM varies. The combination of an FCS (i.e. a nucleic acid binding domain) and a SAM could allow these proteins also to undergo phase separation. In support of this idea, the *C. elegans* SOP-2 protein functions as a PcG protein ^61^, and forms large nuclear bodies ^62^. Although it is not a clear sequence homologue of Ph, SOP-2 consists of an RNA binding motif, an intrinsically disordered region (IDR), and a SAM ^63^. Recently, the IDR of SOP-2 was shown to undergo LLPS *in vitro*, induced by crowding agents or RNA ^41^. Addition of the SAM to the IDR still allowed LLPS, but resulted in formation of smaller condensates that showed lower recovery in FRAP experiments ^41^.

A model for the function of Ph SAM that can reconcile the seemingly different requirements for the SAM and its polymerization activity in different contexts is that Ph SAM drives at least three different states. First, Ph SAM polymerization activity may drive formation of tiny PcG clusters that mediate local repression of transcription simply through cooperative binding interactions. This is consistent with our finding that Ph SAM and its polymerization activity increases the DNA binding affinity of Mini-Ph, at concentrations well below the range where phase separation occurs (Fig. 3). It is also consistent with the dependence of Ph repressive activity when targeted to a reporter gene on Ph SAM polymerization activity ^5^. Second, bridging of nucleosomes mediated by the polymerization interfaces of Ph SAM associated with chromatin bound PRC1 may drive collapse of the chromatin polymer over larger regions of PRC1 bound chromatin ^14,44,64,65^. Indeed, a model of this process could explain the observed effects of overexpressing Ph with the strong ML mutation or wild-type Ph, which increases the number but not size of Ph clusters ^14^. In cases where the local concentration of Ph is very high, Ph may undergo LLPS mediated by multivalent interactions among Ph molecules and between Ph and chromatin (or Ph and RNA), as captured by our *in vitro* assays, and possibly, in the foci observed when Venus-Ph is overexpressed in cells (Fig. 7). Which mechanism dominates in any situation could be modulated by the local concentration of PcG proteins (i.e. how strong a PcG recruitment site is, or the density of recruitment sites). This could be analogous to the distinction between enhancers and super-enhancers, which recruit higher levels of transcription factors and co-factors and where LLPS is believed to occur ^34,66^. There is also no reason at this time to exclude hybrid models ^54^. For example, LLPS could be a mechanism to create biochemical compartments, and within these domains, strict SAM-SAM interactions could establish precise chromatin contacts required for gene repression. LLPS may also represent an extreme and transient state, used to silence large chromatin domains rapidly during development ^12,67^, or as a step in re-establishing gene expression patterns during the cell cycle. All of these possibilities remain to be tested, but the separation of phase separation and polymerization activity revealed by our simple *in vitro* assays may provide a means to do so.

Many proteins with diverse localizations and functions have SAMs. Some SAMs have been shown to polymerize in a concentration dependent manner, while others require additional recruitment mechanisms to induce polymerization. The SAMS of a subset of proteins, including Ets1, Fli1, and p63 ^68^, have not been observed to polymerize. It is therefore possible that phase separation is a property of the SAM that is distinct from polymerization, a hypothesis that is testable by measuring the phase separation activity of proteins with monomeric SAMs.

### Ph SAM and histone ubiquitylation

We find that Ph SAM driven chromatin condensates can enhance PRC1-mediated histone ubiquitylation. We do not know what the mechanism of stimulation of H2A-Ub is. It is unlikely to be concentration of the reaction components in condensates because all of the components (except PRC1ΔPh) are present at saturating concentrations in these reactions. PRC1ΔPh binds chromatin tightly (Kd for 150 bp DNA is <=1 nM ^69^) so that Mini-Ph is also not needed to recruit PRC1ΔPh to chromatin. Although further experiments will be needed to determine the mechanism, the environment of condensates may stimulate steps in the reaction subsequent to substrate binding, which could include the actual ubiquitin transfer or steps affecting processivity ^70^. It has recently been shown that H2A-Ub mediated by PRC1 is stimulated by chromatin compaction ^71^, and that spreading of H2B-Ub along chromatin is facilitated by formation of structured, phase-separated compartments by the ubiquitylation machinery ^72^, which may be relevant to our observations. Formation of protein-chromatin condensates with the heterochromatin protein HP1 alters the conformation of the nucleosome, rendering specific regions of the histone proteins more accessible ^73^. It is possible that nucleosome conformation is also changed in Mini-Ph condensates, and that these changes facilitate histone ubiquitylation. Detailed characterization of chromatin in condensates will be an important future goal.

Stimulation of H2A-Ub is unlikely to be the essential function of the Ph SAM in *Drosophila*, since the modification is not required for PRC1-dependent gene repression *in vivo*, including repression of genes that depend on Ph SAM ^74,75^. However, H2A-Ub is required for full development ^74,75^. *Drosophila* cPRC1 also does not seem to mediate most H2A-Ub in tissue culture cells, and it is likely that another ncPRC1 containing L3(73)Ah, a homologue of mammalian Pcgf3, in place of PSC, is present in these cells ^76^. This also means that in our experiments with nuclear extracts, although we observe PRC1 recruitment to condensates, we cannot be certain that it is responsible for the ubiquitylation activity we observe (Fig. 6).

Histone ubiquitylation by PRC1 has been most intensively studied in mouse embryonic stem cells (mESCs), where systematic analysis of the effect of disrupting PRC1 subunits implicates ncPRC1 (i.e. non PHC-containing) in creation of most H2A-Ub ^7-9^. However, using an artificial tethering system that allows PcG proteins to be reversibly targeted to a reporter gene so that persistent effects on chromatin and gene expression (i.e. memory) can be measured, Moussa et al. ^77^ found that heritable gene repression and propagation of H2A-Ub depend on cPRC1. Recent work indicates a central role for H2A-Ub in PcG-dependent gene regulation in mESCs ^78,79^, in seeming contrast with observations in *Drosophila*; it will be interesting to determine how Ph SAM contributes to H2A-Ub activity in mammals. The ability of Ph SAM to condense chromatin and to promote H2A-Ub could be important for rapidly building PcG chromatin domains, or restoring them at the end of mitosis. H2A-Ub is not detected on mitotic chromosomes in mammalian cells ^80,81^, suggesting it is re-acquired after cells exit mitosis.

Finally, Cbx2, a member of some mammalian canonical (PHC-containing) PRC1s, which has a strong chromatin compacting activity ^82^, has also been shown to form phase separated condensates with chromatin *in vitro*, and to form 1,6-hexanediol-sensitive foci in ES cells ^40,42^. This phase separation activity is mediated by a charged IDR in Cbx2 that is important for the developmental function of Cbx2 ^83^. Further, as shown in Supplementary Fig. 17, Mini-Ph does not form foci in cells, indicating that other sequences in Ph, all of which are predicted to be disordered, can regulate the activity of the Ph SAM. How the activity of Ph SAM is regulated by other sequences in Ph and coordinated with that of other components of PRC1, particularly that of PSC which has a powerful chromatin compacting activity analogous to that of Cbx2 ^84^, is an important question for future study.

## Methods

### Cloning

Cloning of Mini-Ph and the polymerization mutants was described previously ^5^. Mini-PhΔSAM (residues 1291 – 1507) and Mini-PhΔFCS (residues 1397 – 1577) were cloned into a modified pET-3c vector expressing a leader sequence containing a hexahistidine tag followed by a TEV cleavage site. To express Venus-tagged proteins in S2 cells, Ph, Ph-ML, or PhΔSAM were first cloned into a house-modified gateway donor vector and full sequences confirmed. LR recombination was used with pHVW from the DGRC (stock # 1089) to produce the final expression plasmids.

### Protein purification

#### Mini-Ph

His-tagged Mini-Ph, Mini-Ph-EH, and Mini-Ph-ML were expressed in Rosetta (DE3) *E. coli*. Cultures were grown at 37°C to an OD of 0.8-1.0, and then shifted to 15°C for overnight induction with 1mM IPTG. Cells were pelleted, flash frozen, and stored at −80°C. Cells were resuspended in 2 ml/g lysis buffer (50 mM Tris, pH 8.5, 200 mM NaCl, 10 mM β-ME, 100 µM ZnCl_2_, 0.2 mM PMSF, 0.5 mM Benzamidine). Cells were incubated on ice for 10 min, flash frozen in liquid nitrogen, thawed at 37°C, and sonicated 6*30 sec. at 30% intensity. Freeze-thaw and sonication were repeated, and the lysate centrifuged for 1 hour at 100,000*g and 4°C. Cleared lysate was sonicated 6*30” at 40% intensity, and filtered through a 22 µm filter. Lysate (from 1 L) was applied to a 1 ml His-Trap column using an AKTA FPLC, and eluted with a gradient of imidazole (from 10-300 mM) in lysis buffer. Fractions with Mini-Ph were dialyzed overnight against 1 L of 20 mM Tris, pH 8.5, 50 mM NaCl, 100 µM ZnCl_2_, and 10 mM β-ME. Dialyzed fractions were centrifuged for 10 min. at 20,800*g, and loaded on a 1 ml HiTrapQ-HP column and eluted with a gradient from 50 mM to 1 M NaCl in binding buffer. Fractions were pooled and dialyzed overnight into 20 mM Tris, pH8, 50 mM NaCl, 10 µM ZnCl_2_, 1 mM βME, aliquotted and stored at −80. In some cases, Mini-Ph was further purified by size exclusion chromatography on a Superose 12 size exclusion column.

#### Mini-PhΔSAM and Mini-PhΔFCS

Both proteins were expressed in BL21 (DE3) Gold cells pre-transformed with the pRARE plasmid. The transformed cells were grown at 37°C in LB media to an OD_600_ of ∼0.7 – 0.8 and induced overnight at 15°C. Cells harvested from 1 L of culture were resuspended with 10 ml of lysis buffer (50 mM Tris pH 8.0, 200 mM NaCl, 5 mM βME, 30 mM imidazole pH 7.5, 1 mM PMSF) and lysed by sonication. The soluble lysates were introduced onto am Ni-NTA column, washed with lysis buffer (without PMSF), and bound proteins eluted using 300 mM imidazole, 200 mM NaCl, 5 mM βME. The leader sequence was cleaved using TEV protease, and the cleaved sequence and uncleaved proteins removed by passing through a Ni-NTA column. Further purification was performed using a HiTrap Q-HP column. Fractions containing protein were pooled, buffer exchanged into 50 mM Tris pH 8.0, 100 mM NaCl, 5 mM βME, and concentrated. Mini-PhΔSAM was further purified on a Superdex 200 size exclusion column in 50 mM Tris pH 8.0, 100 mM NaCl, 5 mM βME. Purified, concentrated proteins were stored at −80°C

#### E1, E2, and His-Ub

The following plasmids were used: human 6X-His-UBA1 (E1) (pET21d-Ube1, addgene #34965), Human UbcH5c (E2) (pET28a-UbcH5c, addgene # 12643), 6XHis-Ubiquitin (pET15b-His-Ub) (kind gift of B. Schulman). Proteins were expressed in *E*.*coli* and purified essentially as described ^85,86^. His-Ube1 was purified by Ni-NTA affinity followed by Superdex 200 chromatography ^85^. UbcH5c was purified on a HiTrap SP-XL column followed by Superdex 200 ^86^. 6X-His-Ub was purified by Ni-NTA chromatography.

#### Histone purification

*Xenopus laevis* histones, including H2B-122C mutant were expressed in and purified from *E. coli*, as described ^87,88^. All experiments were carried out with histone H3 with Cys110 (the only cysteine natively present in the histones) mutated to Ala.

Fluorescent labeling of histone H2A with NHS-Cy3 was carried out under conditions favouring labeling of the N-terminal amine. Lyophilized H2A was resuspended in labeling buffer (20 mM Hepes, pH 6.2, 7 M Guanidium-HCl, 5 mM EDTA) to a concentration of 0.1 mM. NHS-Cy3 stock (in DMF) was added to a final ratio of 0.5:1 (dye to histone) and incubated at room temp. for 90 min. Free dye was removed with Amicon concentrators, after diluting with labeling buffer without Guanidium to reduce the Gu-HCl concentration to 6 M. In some cases, Zeba columns were used instead to remove free dye. To label H2B-122C with maleimide-Alexa 647, lyophilized histone was reconstituted in denaturing labeling buffer (20 mM Tris-HCl, pH 7.0, 7 M guanidium HCl, 5 mM EDTA) to a final concentration of 0.1 mM followed by treatment with a 100-fold excess of TCEP for 30 minutes. Maleimide-Alexa 647 was added to a final ratio of 3:1 (dye:histone) and incubated for 3 hours at room temp. The labeling reaction was quenched with β-ME (final concentration 80 mM), and free dye removed as above. Octamer reconstitutions and purification on a Superdex 200 size exclusion column were carried out as described ^87,88^. Concentrated octamers were dialyzed into octamer refolding buffer (2 M NaCl, 10 mM Tris, pH 7.5, 1 mM EDTA, 5 mM β-ME) with 50% glycerol and stored at −80°C.

#### PRC1ΔPh

PRC1ΔPh was purified from nuclear extracts of baculovirus-infected Sf9 cells essentially as described previously ^69^, with the following changes. Nuclear extracts were prepared from Sf9 cells infected with viruses for the three subunits (Flag-PSC, Pc, dRING) but nuclei were purified through a sucrose cushion prior to nuclear extraction. During the purification, the 2 M KCl wash in the published protocol was replaced with a wash consisting of BC2000N + 1 M Urea (20 mM Hepes, pH 7.9, 2 0.4 mM EDTA, 2 M KCl, 1 M deionized urea, 0.05% NP40, no glycerol). Additionally, prior to eluting the protein, anti-FLAG beads were incubated 3-5 volumes of BC300N with 4 mM ATP + 4 mM MgCl_2_ for 30 min. at room temperature. This step reduces the amount of HSC-70 that co-purifies with PRC1ΔPh. Protein was eluted with 0.4 mg/ml FLAG in BC300 without NP40, concentrated to ∼1 mg/ml and stored in BC300N (20 mM Hepes, pH 7.9, 300 mM KCl, 0.2 mM EDTA, 20% glycerol, 0.05% NP40).

### Fluorescent labelling and acetylation of Mini-Ph and other proteins

To fluorescently label proteins, NHS-ester-Cy3 or Alexa-647 were used to randomly label lysines. A Zeba column (Thermo Fisher) was used to buffer exchange the protein into 20 mM Hepes, pH 7.9, 200 mM NaCl for Mini-Ph, or BC300N for proteins expressed in Sf9 cells; labeling was carried out with a 0.5:1 (dye:protein) ratio for 15 min. at room temp. Labelling was quenched by addition of Lysine to 10 mM. Free dye was removed using two Zeba columns, which were equilibrated in the labeling buffer with 200 mM NaCl. Labelled protein was mixed with unlabelled at a ratio of 1 to 25 for imaging experiments. Acetylation of Mini-Ph was carried out exactly as for fluorescent labelling except that a ratio of 8:1 sulfo-NHS-acetate:lysine residues in Mini-Ph was used and labeling was carried out for 1 hour at room temp.

### Preparation of Nuclear Extracts from *Drosophila* S2R+ cells

S2R+ cells were grown in M3-BYPE media with 10% FBS. 20*15 cm dishes were used to prepare nuclear extracts as described ^89^, except that nuclei were purified through a sucrose cushion prior to extraction. Cells lysed in hypotonic buffer were layered over two volumes of 30% sucrose in hypotonic buffer, and centrifuged 18’ @ 1400*g. Nuclei were washed once in hypotonic buffer, and extracted as described. The high salt extraction buffer was 1.2 M KCl, and extracts were not dialyzed. To use the extracts to treat condensates, up to 100 µl of extract was buffer exchanged into 20 mM Tris, pH 8, 50 mM NaCl using a Zeba column. Extracts were centrifuged 2’ @ 20,000*g and incubated for 15’ on ice with 60% volume of Q-sepharose. Extracts were spun through an empty column (2’ @ 10,000*g), and then centrifuged 2’ @20,000 *g. All procedures were carried out on ice or at 4°C and contained protease inhibitors and 0.4X PhosStop phosphatase inhibitor.

### Chromatin preparation

Most experiments were carried out with the plasmid p5S*8, which contains 5 blocks of 8-5S nucleosome positioning sequences (repeat length 208 base pairs). Plasmids were assembled by salt gradient dialysis as described ^90^. Chromatin was finally dialyzed into HEN (10 mM Hepes, pH 7.9, 0.25 mM EDTA, 10 mM NaCl) buffer and stored at 4°C. To measure chromatin assembly, 100 ng of each assembly was digested overnight with 10 U of EcoRI in NEB buffer 2.1, and loaded on a 0.5X TBE, 5% acrylamide native gel. Gels were stained with Ethidium bromide and imaged on a Typhoon imager. For quantification, the nucleosomal signal is multiplied by 2.5 to account for the quenching effect of bound protein on ethidium bromide ^91^. For Micrococcal nuclease analysis, 800-1000 ng of chromatin was diluted into 40 µl of the following buffer: 12 mM Hepes, pH 7.9, 0.12 mM EDTA, 60 mM KCl, 2 mM MgCl_2_ and split into 4 tubes. Micrococcal nuclease (Sigma, #N3755) (0.5 U/µl in 50 mM Tris, pH 8, 0.05 mM CaCl_2_, 50% glycerol) was diluted 1:18, 1:54, 1:162, and 1:486 in MNase dilution buffer (50 mM Tris, pH8, 10 mM NaCl, 126 mM CaCl_2_, 5% glycerol). 1 µl of each dilution was used to digest chromatin for 7 min. at room temp. Reactions were stopped with DSB-PK (10X stock: 50 mM Tris, pH 8.0, 0.1 M EDTA, 1% SDS, 25% glycerol + 10 mg/ml Proteinase K), digested overnight at 50°C, and analyzed on 1X TBE-1.5% agarose (SeaKem) gels which were stained with Ethidium bromide and imaged on a Typhoon Imager.

### Phase separation assays

Proteins and templates were routinely centrifuged full speed in a microfuge for 2-5 min. at 4°C to remove aggregates before setting up phase separation assays. For phase separation assays, reactions (10-20µl) were assembled in a 384-well glass-bottom imaging dish (SensoPlate, Greiner Bio-One). Wells were not pre-treated; pre-coating with BSA did not influence phase separation by Mini-Ph. Phase separation was initiated by addition of the protein or the DNA, and mixing the reaction by gentle pipetting, with care taken not to introduce air. Reactions were incubated in the dark for 15 min. or up to several hours. For reactions where YOYO-1 (Thermo Fisher) was used, it was added at the beginning of the reaction to a final dilution of 1:3000. Typical reaction conditions are 50 mM NaCl or 50 mM KCl, 20 mM Tris pH8. Reactions were set up on ice, and transferred to room temp. for 15 min. Turbidity measurements were made in duplicate using a NanoDrop spectrophotometer. Phase separated condensates were pelleted by centrifugation at 14,000*g for 2 min. at 4°C, and supernatants removed to fresh tubes. Pellets were resuspended in 12 µl 1.5X SDS-Sample buffer, and 6X SDS-Sample Buffer was added to the supernatant. 10% of the pellet and supernatant were removed and digested in DSB-PK for 2 hours at 50°C for DNA analysis. The remainder of the sample was boiled and analyzed by SDS-PAGE.

### Imaging of condensates

All images were collected on a Zeiss microscope, equipped with a Yokogawa CSU-1 spinning disc confocal head. Zen 2012 software was used for image acquisition with a 63X oil objective, or a 100X oil objective (for movies and FRAP) and evolve EMCCD camera from Photometrics. The excitation wave length for YOYO/Venus, Cy3/RFP and Alexa 647 were 488, 561 and 639 nm respectively

### Measuring nucleosome concentration in condensates

Images were collected at 25% laser power, 200 msec exposure for buffer, chromatin alone, a titration of labelled histone octamers (in octamer refolding buffer, which contains 2M NaCl, and in which histone octamers remain assembled), and Mini-Ph chromatin condensates. Histones are the same histones used to prepare the chromatin; 43% of the histone octamers are labelled (measured both using the NanoDrop and by loading histones and free dye on SDS-PAGE gels), corresponding to a 21.5% labeling efficiency on H2A (since there are two copies of H2A in each octamer). Image J “measure” was used to measure the mean grey intensity for each of 9 images for each point. Images were manually checked and images with bright artifacts removed, although these had little impact on the measured intensities. A linear regression was fit to the calibration curve and used to convert measured intensities to nucleosome concentrations. To measure intensities in condensates, Image J was used to threshold the images (AutoThreshold-->Li); Analyze Particles was used to measure the mean grey intensity in each thresholded structure. Particle size was set as 100-infinity pixels. The mean grey intensity from the buffer image was subtracted from all measurements, which were converted to nucleosome concentrations using the calibration curve.

### FRAP

FRAP experiments were carried out with Alexa-647 labelled Mini-Ph or Mini-Ph EH. Bleaching was done with a 595 nm laser, for 1500 msec. This effectively bleaches both Alexa-647 Mini-Ph, and Cy3-H2A, although we were only able to record FRAP images from one channel. Two pre-bleach images were collected, followed by an image every 5 or 10 sec. All FRAP analysis of Mini-Ph was done by bleaching single complete structures. Images were analyzed in Image J (Fiji). An ROI was selected for the bleach area, background, and a non-bleached structure. Background subtracted, normalized data were fit with a double exponential fit using GraphPad Prism 8.

Y=Y0+ SpanFast*(1-exp(-KFast*X)) + SpanSlow*(1-exp(-KSlow*X)). We excluded data sets that could not be fit, and obvious technical artifacts (e.g. if a drop fuses with the bleached condensate during the experiment).

### Image analysis of condensates

Images for display were prepared using Zen2 (blue edition). For quantification, images were exported as TIFs from Zen (original data). ImageJ (Fiji) was used to threshold the images (Li algorithm); thresholds were manually checked and images with too few structures to threshold were removed. Areas of thresholded structures were measured using ImageJ (“Analyze Particles”, size=10-infinity pixel), and intensities using Analyze Particles. For colocalization analysis, the GDSC-->Colocalization-->Particle Overlap was used. Masks were created in the Alexa 647 (Mini-Ph) and Cy3 (chromatin) channels, and overlap of Cy3 with Mini-Ph structures measured.

Movies were created from .czi files in ImageJ (Fiji). Movies were saved as .avi files at 1, 2, or 3 frames per second, and using PNG compression.

### Filter binding

Filter binding was carried out as described ^69,92^. Briefly, a 150 bp internally labelled DNA probe was prepared by PCR and gel purified. The probe was used at 0.02 nM. Reaction conditions were 60 mM KCl, 12 mM Hepes, pH7.9, 0.24 mM EDTA, 4% glycerol, in a 20 µl volume. Proteins were centrifuged 2 min. at full speed in a microfuge before preparing the dilution series. Binding reactions were incubated 1 hour at room temperature. Hybond-XL was used as the bottom membrane (binds DNA), and was pre-equilibrated in 0.4 M Tris, pH 8. Nitrocellulose was used as the top membrane (binds protein + DNA), and was pre-treated with 0.4 M KOH for 10 min., neutralized by washing through several changes of Milli-Q water, and equilibrated for at least 1 hour in binding buffer. Filters were assembled in a 48-well slot-blot apparatus, and each well washed with 100 µl binding buffer. The vacuum was turned off, and reactions loaded on the filters. Slots were immediately washed with 2*100 µl binding buffer. Filters were air dried, exposed to a phosphoimager screen, and scanned on a Typhoon. ImageQuant was used to quantify top (bound) and bottom (unbound) filters, and fraction bound calculated in Excel. Curve fitting was done in GraphPad Prism 8, using the following equation:

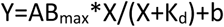

### Protein footprinting assay

The acetylation footprinting assay is described in detail in Kang et al. ^6^. Phase separation reactions were directly scaled up to use 4 µg of protein for each sample. Condensates were allowed to form at room temp. for 15 min.; an aliquot of each sample was removed to confirm phase separation using microscopy. Sulfo-NHS-acetate was dissolved immediately before use, and added to a final concentration of 0.5 mM. An aliquot of each sample was removed to monitor phase separation by microscopy, and reactions were stopped after 15 min. by addition of Trifluoroacetic acid to a final concentration of 1%. For Mini-Ph EH, acetylation of condensates was restricted to 5 min. because these condensates dissolved rapidly on exposure to Sulfo-NHS-acetate. We therefore analyzed Mini-Ph EH alone, and bound to DNA (16X DNA, Fig. 4) after both 5 and 15 min. of acetylation. Samples were TCA precipitated, denatured with 8 M urea, reduced with DTT (45 mM final concentration), treated with a final concentration of 10 mM Iodoacetamide, and diluted 1:2 with H_2_O before treating with Propionic anhydride twice. Samples were dried, treated with Propionic anhydride again, dried, resuspended and digested sequentially with Trypsin and Chymotrypsin. Samples were purified with a ZipTip and analyzed by LC-MS/MS on an Orbitrap-Fusion mass spectrometer.

Mass Spectrometry data were analyzed using Maxquant (v 1.6.10.43) with Acetyl(K) and Propionylation(K) as variable modifications. 10 missed cleavages were allowed since lysine modification will block trypsin digest. All data files were analyzed together, with the “match between runs” option. Intensities for identified Acetyl and Propionyl sites were used for quantification. Accessibility was calculated (in Excel) as (intensity acetylated)/(intensity acetylated+intensity prop +0.5) for each site. To compare accessibility between samples, GraphPad Prism 8 was used to conduct student’s t-test, assuming equal variance across samples, and with the Holm-Sidak method of correction for multiple comparisons, with alpha=0.05 (unpaired, 2-tailed test). Heat maps were prepared from averaged accessibilities using Morpheus (https://software.broadinstitute.org/morpheus).

### Analysis of condensates after incubation in nuclear extracts

Phase separation reactions were set up in 40 µl with 80 nM nucleosomes, 7.5 µM Mini-Ph, in 20 mM Tris pH 8.0 and 50 mM NaCl. After incubating 10 min. at room temp., 12 µl of nuclear extracts were added, and reactions mixed by gently pipetting up and down. 7.5 µl were removed and diluted to 10 µl for imaging, and 7.5 µl mixed with the uibiquitylation machinery to assay histone ubiquitylation. After 60 min. of total incubation, samples were pelleted by centrifugation for 2 min. at 2500*g, 4°C. Supernatants were removed and SDS-sample buffer added to 1X. Pellets were resuspended in 2X SDS sample buffer. 2 µl of each pellet and supernatant were removed and digested with Proteinase K for at least 1 hour at 55°C before analysis on 1.2% agarose, 1X TAE gels, which were stained with SYBR Gold to visualize nucleic acids. The remainder of the samples were boiled and loaded on 8% SDS-PAGE gels, transferred to nitrocellulose, and used for Western blotting. Membranes were blocked with 5% nonfat dry milk in PBST (PBS + 0.3% Tween-20), and incubated with primary antibodies diluted in 5% milk-PBST overnight at 4°C. Membranes were washed 3*10 min. in PBST, incubated in secondary antibody diluted in 5% milk-PBST for 1-2 hours, washed 3*10 min. in PBST, and visualized using a Li-Cor Odyssey imaging system. Image J (Fiji) was used to quantify band intensities.

### Histone Ubiquitylation assays

For ubiquitylation assays, 125 ng chromatin per 5 µl was pre-incubated with 5 µM Mini-Ph (or buffer) for 15 min. at room temp. to induce phase separation, followed by addition of the ubiquitylation machinery and PRC1ΔPh. Final reaction conditions are 40 nM nucleosomes, 20 mM Hepes, pH 7.9, 0.25 mM MgCl_2_, 0.25 mM ATP, 0.6 mM DTT, 60 mM KCl, 25mM NaCl, 700 nM E1, 800 nM E2, and 500 ng Ub. Titrations of the E1, E2, and His-Ub indicate that none are limiting under these conditions. Reactions were further incubated for 45 min. at room temp. Aliquots were removed for imaging, and the remainder of the reaction stopped by addition of SDS-Sample buffer. Boiled samples were loaded on 16% SDS-PAGE gels, which were scanned for Cy3 to detect H2A, and then stained with SYPRO Ruby. Histone ubiquitylation assays in nuclear extracts were carried out under the same conditions except that the pre-incubation of chromatin with Mini-Ph was 10 min., nuclear extracts were added just before the ubiquitylation components, and reactions were incubated for 80 min. at room temp.

### Cell culture

Wild type S2 cells (from Expression Systems) and S2 cell lines harbouring stable Ph or Ph-ML ^14^transgenes were grown in suspension in ESF-921 media with 5% FBS. Protein expression was induced with 0.5 µM CuSO_4_ for 4 days. For whole cell extracts, cells were resuspended in 2X-SDS sample buffer and boiled. For histone extraction, we followed the protocol of Abcam (https://www.abcam.com/protocols/histone-extraction-protocol-for-western-blot); HDAC inhibitors were not included. Western blots were carried out as described above, and ImageQuant was used to quantify bands.

### Live cell imaging

For live cell imaging, S2 (Fig. 7), or S2R+ (Supplementary Fig. 17) cells were plated at 10^6^ cells per well in 6-well plates the night before transfection. Transfection was carried out using Trans-IT lipid (Mirus), according to the manufacturer’s protocol. 2 µg of each Venus-Ph construct was used along with 0.5 µg of pAct5C-H2A-RFP ^93^. One to two days after transfection, cells were replated on ConA-coated imaging dishes (Ibidi). Heat shock was for 8 min. (S2R+) or 12 min. (S2) at 37°C, and cells were analyzed within 24 hours of protein induction. Confocal stacks of thick slices (3 µm) were collected on the spinning disc microscope described above using the 63X objective to capture foci throughout the cell.

### Image analysis of live cells

The .czi files of image stacks were opened in Image J (Fiji), converted to maximum intensity projections, and the channels split. The red channel (H2A-RFP) was used to segment nuclei as follows. Images were thresholded with the Li algorithm, followed by removing outliers less than 5 pixels, and 3 rounds of erosion. Thresholded images were converted to masks, processed with a watershed algorithm, and “Analyze Particles” used with a size threshold of 200-inifinity pixels to select nuclei. The green channel (YFP fusion proteins) was then processed with “Find maxima” with the following parameters: Prominence: 20000; strict; exclude edge maxima; output type: single points. The nuclei selected from the red channel were used as ROIs, and the # maxima per ROI (i.e. # foci/nucleus) obtained using Measure in the ROI tool, followed by dividing the raw integrated density by 255. This entire pipeline is explained here: https://microscopy.duke.edu/guides/count-nuclear-foci-ImageJ. To compare the # foci per cell, cells with zero foci were excluded; since Venus-PhΔSAM does not form foci, the majority of cells were excluded.

### Data Availability

Mass spectrometry raw files will be uploaded to MassIVE. The Source Data file includes data for FRAP traces (Fig. 2, Supplementary Fig. 14) and MaxQuant output (intensities) for acetylation footprinting experiments (Fig. 4), filter binding data (Fig. 3C), nucleosome and condensate measurements (Fig. 3H, I, J), western blot quantification (Fig. 5I, 7C, D), ubiquitylation activity (Fig. 6E), foci measurements (Fig. 7H). All other raw data are available on reasonable request.

## Supplemental information

**Supplementary Figures 1-18**

**Supplemental Movie Legends 1-3**

**Supplementary Table 1**

**Supplementary Figure 1.**
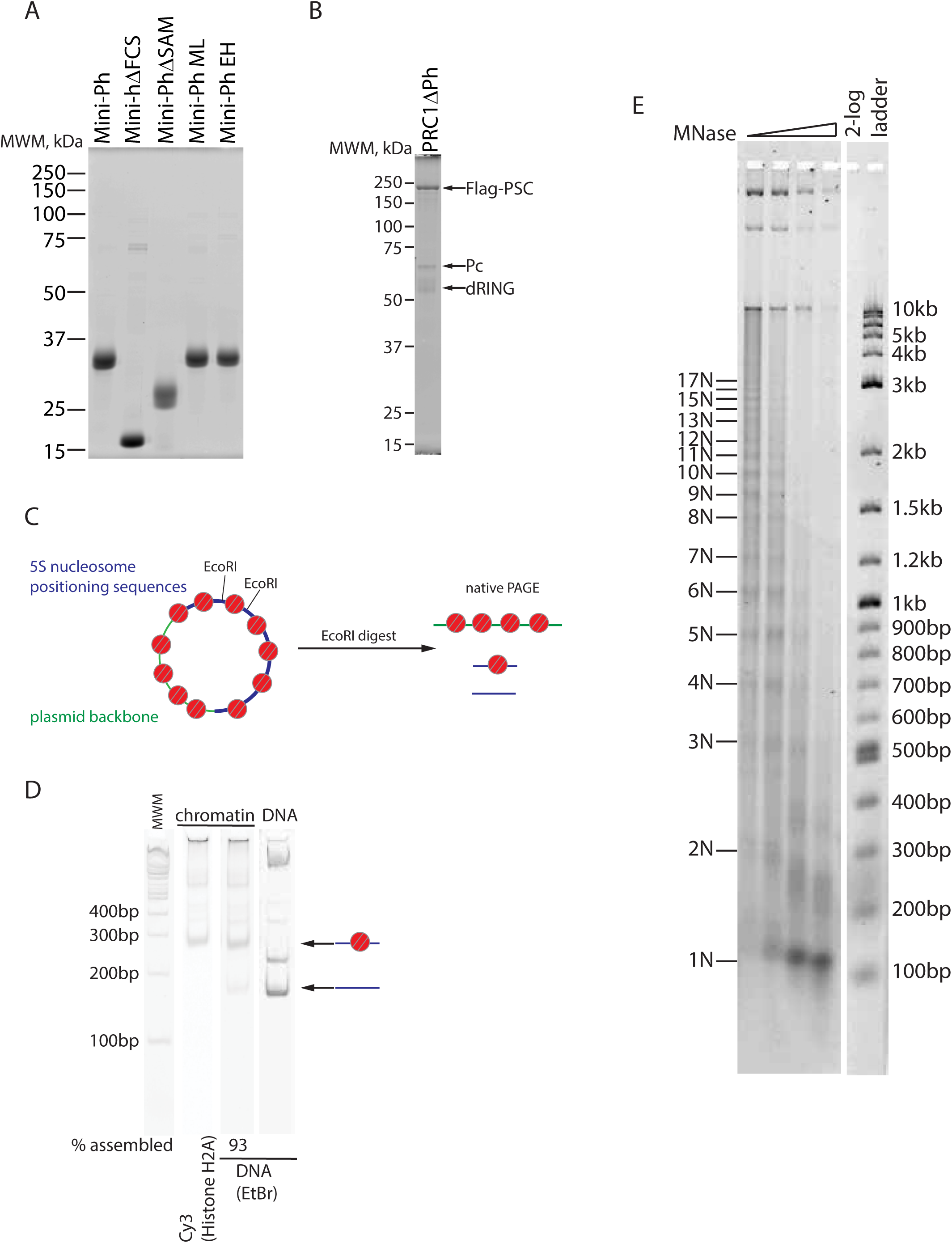
Proteins and Chromatin used in these experiments. A. SYPRO Ruby stained SDS-PAGE of Mini-Ph and its derivatives that were prepared in *E. coli*. Equimolar amounts of each protein were loaded. B. Ruby stained gel of 3-subunit PRC1 consisting of Flag-PSC, Pc, and dRING (PRC1ΔPh). C. Schematic of plasmid used for chromatin assembly. The plasmid used for most experiments consists of 40*5S nucleosome positioning sequences (208 base pair repeat) (blue), and the plasmid backbone (green). Each 5S sequence is flanked by EcoRI sites. To estimate nucleosome assembly, chromatinized plasmids are digested with EcoRI, and the naked and nucleosomal 5S repeats resolved by native PAGE. D. Representative gel of EcoRI digest. Left shows DNA stain used to quantify naked and nucleosomal repeats, and right side shows the Cy3 label on histone H2A. E. Micrococcal nuclease analysis of chromatin used for condensate formation. Numbers on the right indicate fragments representing nucleosomal increments.

**Supplementary Figure 2.**
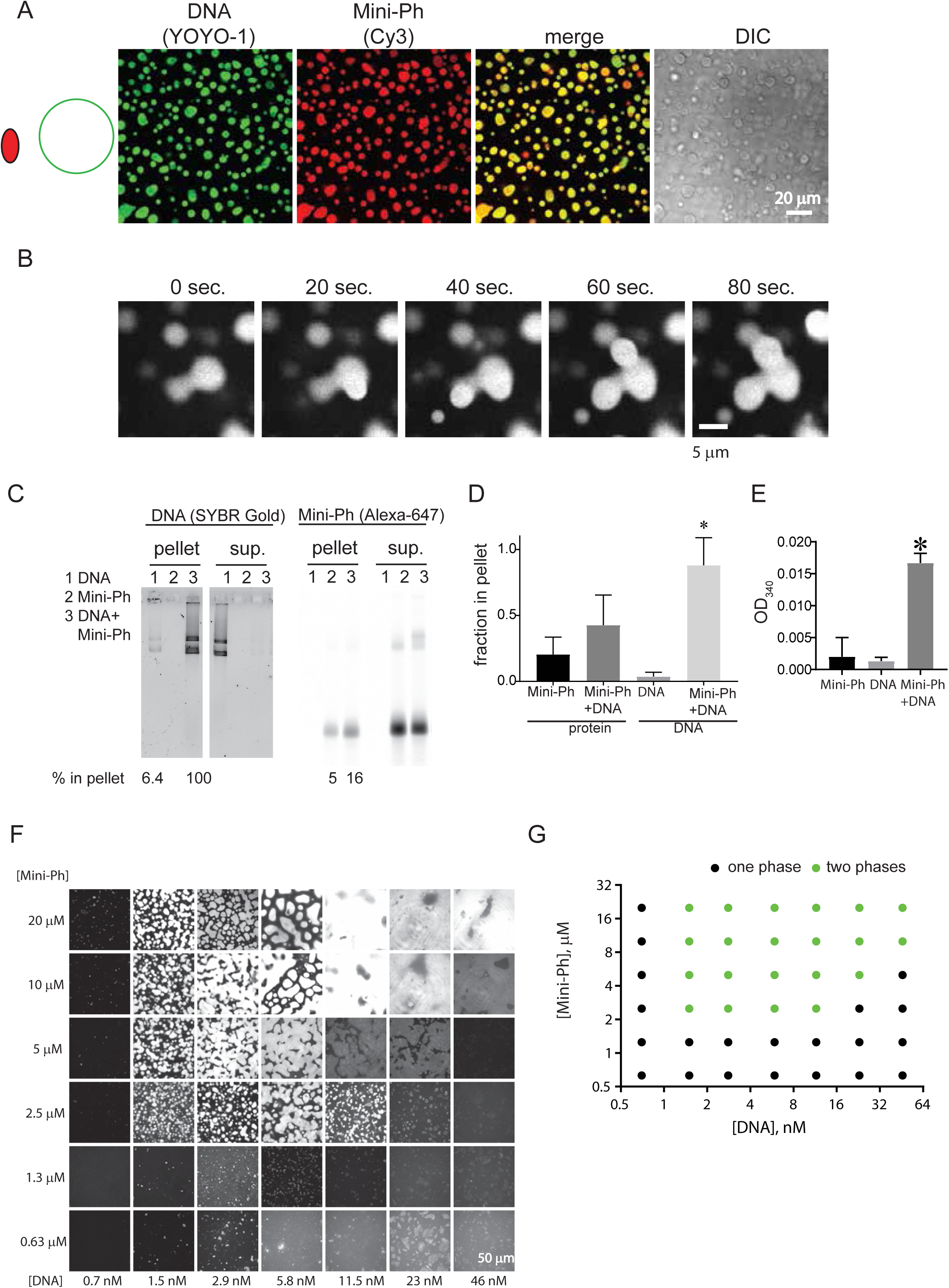
Mini-Ph forms phase separated condensates with DNA. A. Mini-Ph forms phase separated condensates with DNA. B. Time-lapse of droplet fusion of Mini-Ph-chromatin condensates, visualized with Alexa 647-labelled Mini-Ph. C-E. Mini-Ph-DNA condensates can be pelleted by brief centrifugation. Representative gels (C) of DNA (left) and protein (fluorescent scan of Alexa 647-labelled Mini-Ph). Condensates were allowed to form for 15 min. at room temperature. D. Quantification of pelleting experiments. E. Mini-Ph-DNA condensates increase turbidity measured by OD_340_. * indicates p<0.05, student’s t-test. F. Matrix of Mini-Ph and DNA. G. Graph of one-phase versus two-phase regions of the Mini-Ph-DNA matrix. Note that high concentrations of DNA disrupt condensate formation (e.g. 46 nM DNA with 5 μM Mini-Ph). [DNA] refers to the concentration of plasmids.

**Supplementary Figure 3.**
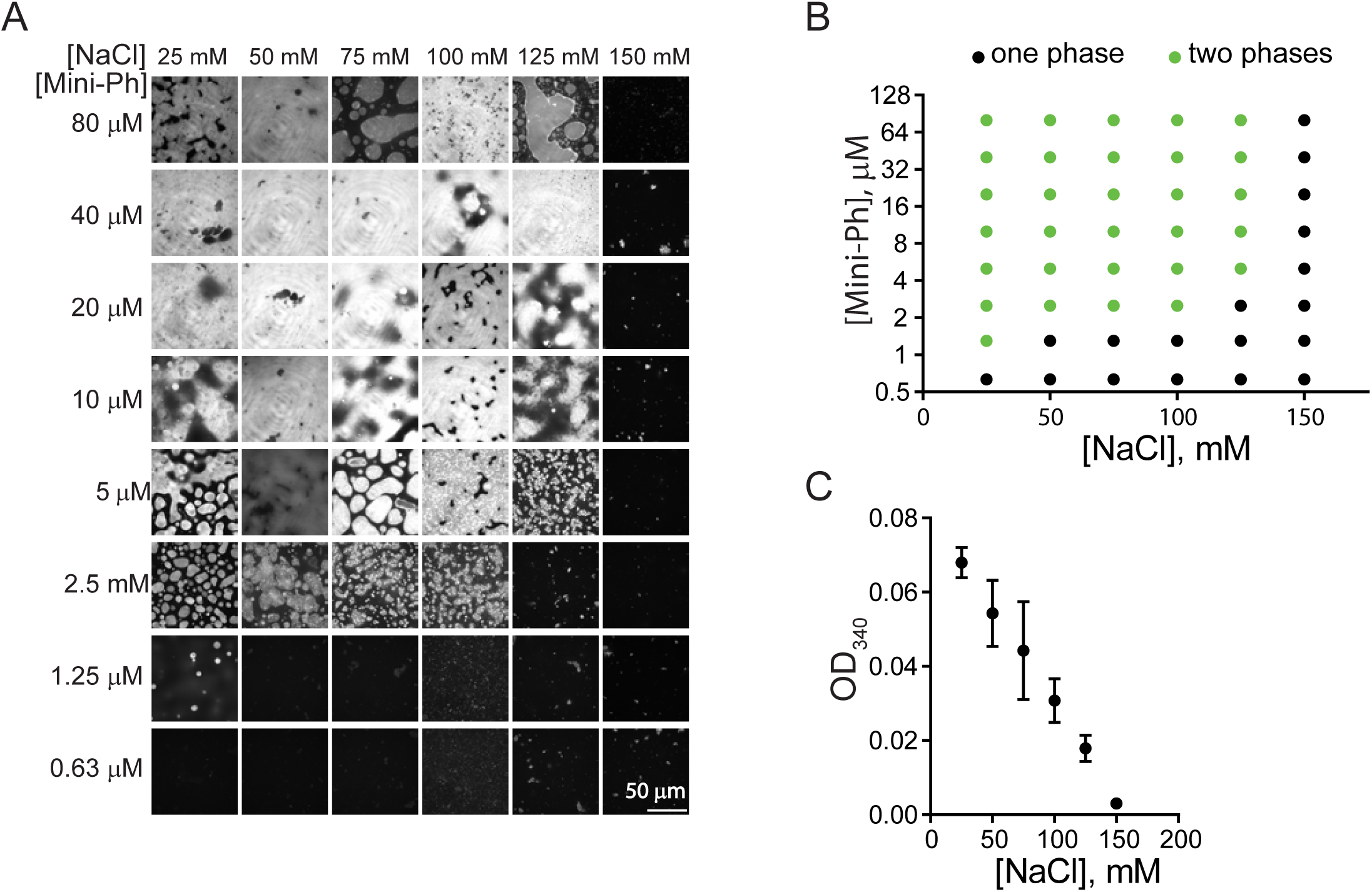
Mini-Ph-DNA condensates are sensitive to NaCl concentration. A, B. Matrix of of Mini-Ph-DNA condensates and across different concentrations of NaCl. A shows representative images, and B the graph of one-phase versus two-phase points. C. Increasing [NaCl] disrupts condensates as measured by OD_340_. Mean +/- SD of three titrations.

**Supplementary Figure 4.**
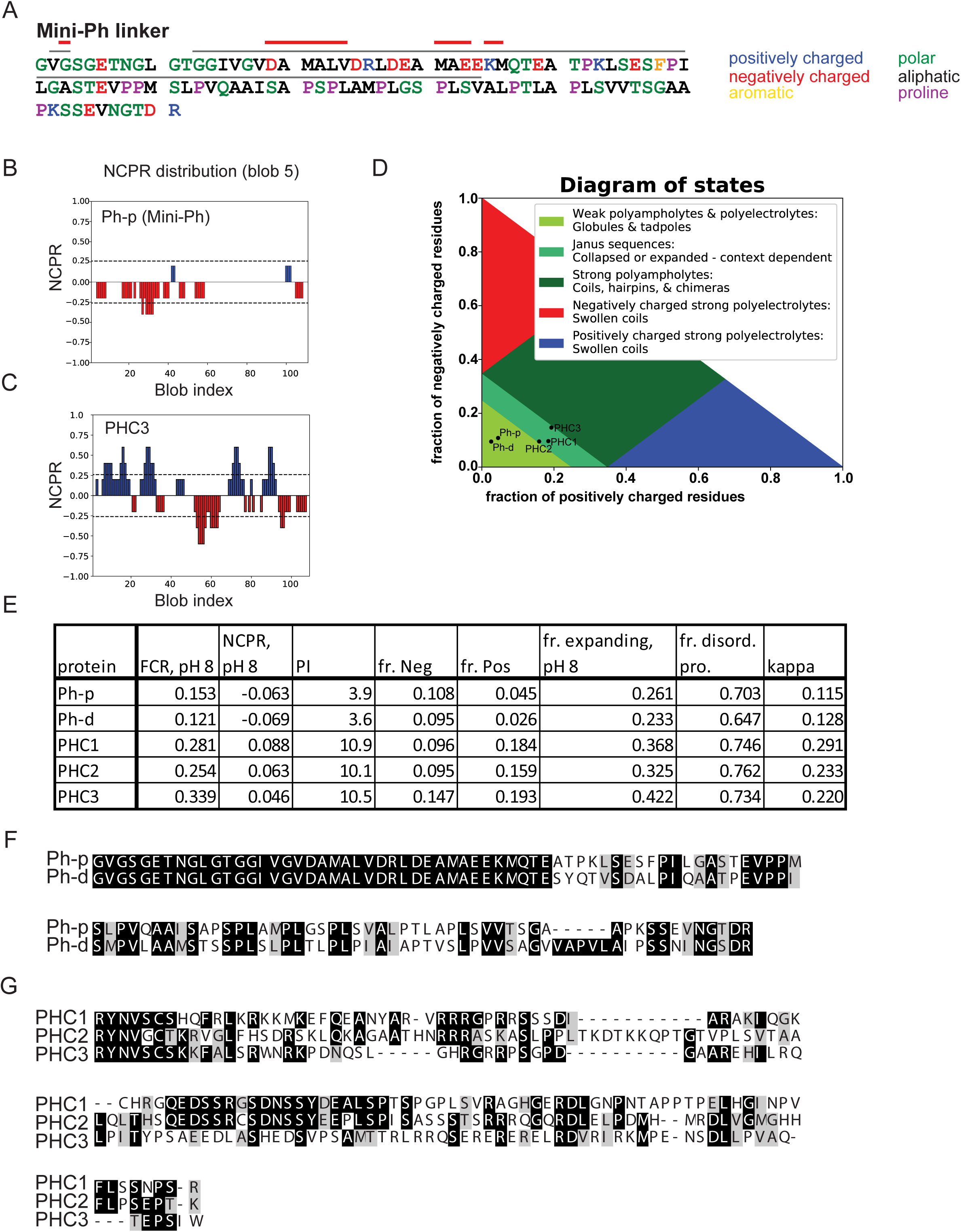
Sequence properties of Mini Ph linker. A. Colour coded Mini-Ph linker sequence. Gray and red lines indicate linker residues that may interact with Ph SAM based on NMR results with the linker and the SAM in trans. Red line indicates residues that had altered NMR signals when mixed with 0.4 molar equivalents of Ph SAM, and gray lines those with altered signals with 1.6 molar equivalents of Ph SAM. See Robinson et al. (2012, JBC) for details. B. Plot of net charge per residue (NCPR, pH 8.0) of Mini-Ph linker indicating charge distribution in the linker. C. Plot of NCPR (pH 8.0) for PHC3, indicating distinct charge patterning compared with Ph-p. D. Das-Pappu diagram of states for both *Drosophila* and all three human Ph linkers, showing that the *Drosophila* and human linkers have distinct properties. E. Summary of properties of Ph linkers calculated with http://pappulab.github.io/localCIDER/. See also Supplementary Table 1 for all sequences and parameters (FCR=fraction charged residues). F. Alignment of Ph-p and Ph-d linkers. G. Alignment of PHC1-3 (human Ph homologues) linkers.

**Supplementary Figure 5.**
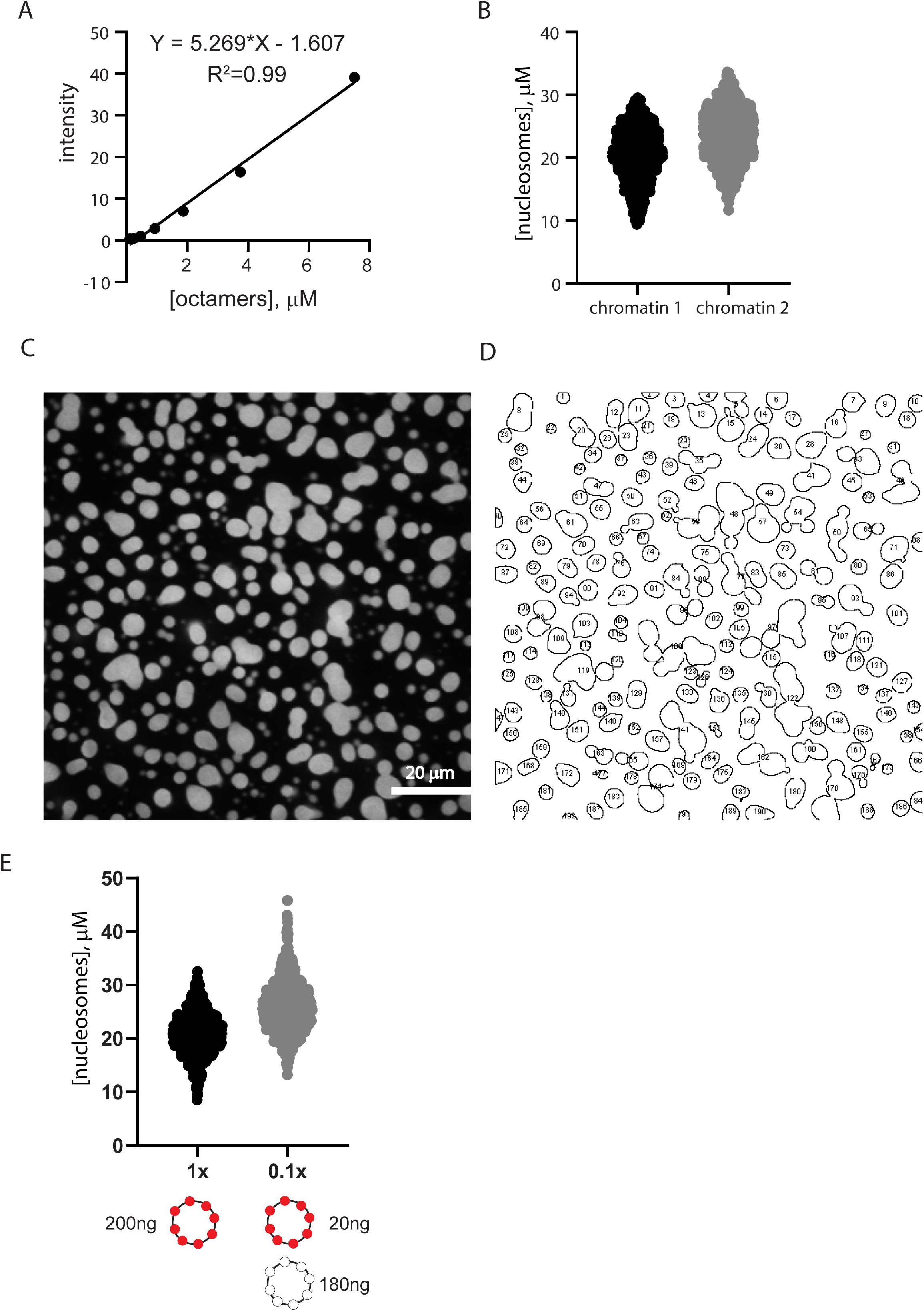
Quantification of nucleosome concentration in condensates. A. Standard curve prepared from serial dilution of Cy3-labelled (on H2A) histone octamers. Buffer alone was used to collect a background intensity, which was subtracted, from all images. B. [nucleosomes] in individual structures measured from two reactions carried out with different chromatin assemblies. Chromatin 2 was assembled at higher nucleosome density, presumably explaining the higher nucleosome concentration in condensates. C. Example image used for quantification of nucleosome concentration. D. Outlines of thresholded structures selected for quantification for the image shown in C. E. Graph of experiments comparing concentrations of structures measured with a 1:9 mixture of Cy3-labelled (0.1X) to unlabelled (1X) chromatin versus all Cy3-labelled. The calculated concentrations are similar (21+/-4 µM and 26+/-5 µM) in both reactions, suggesting the measurements in the dense phase with all labelled chromatin are in the linear range.

**Supplementary Figure 6.**
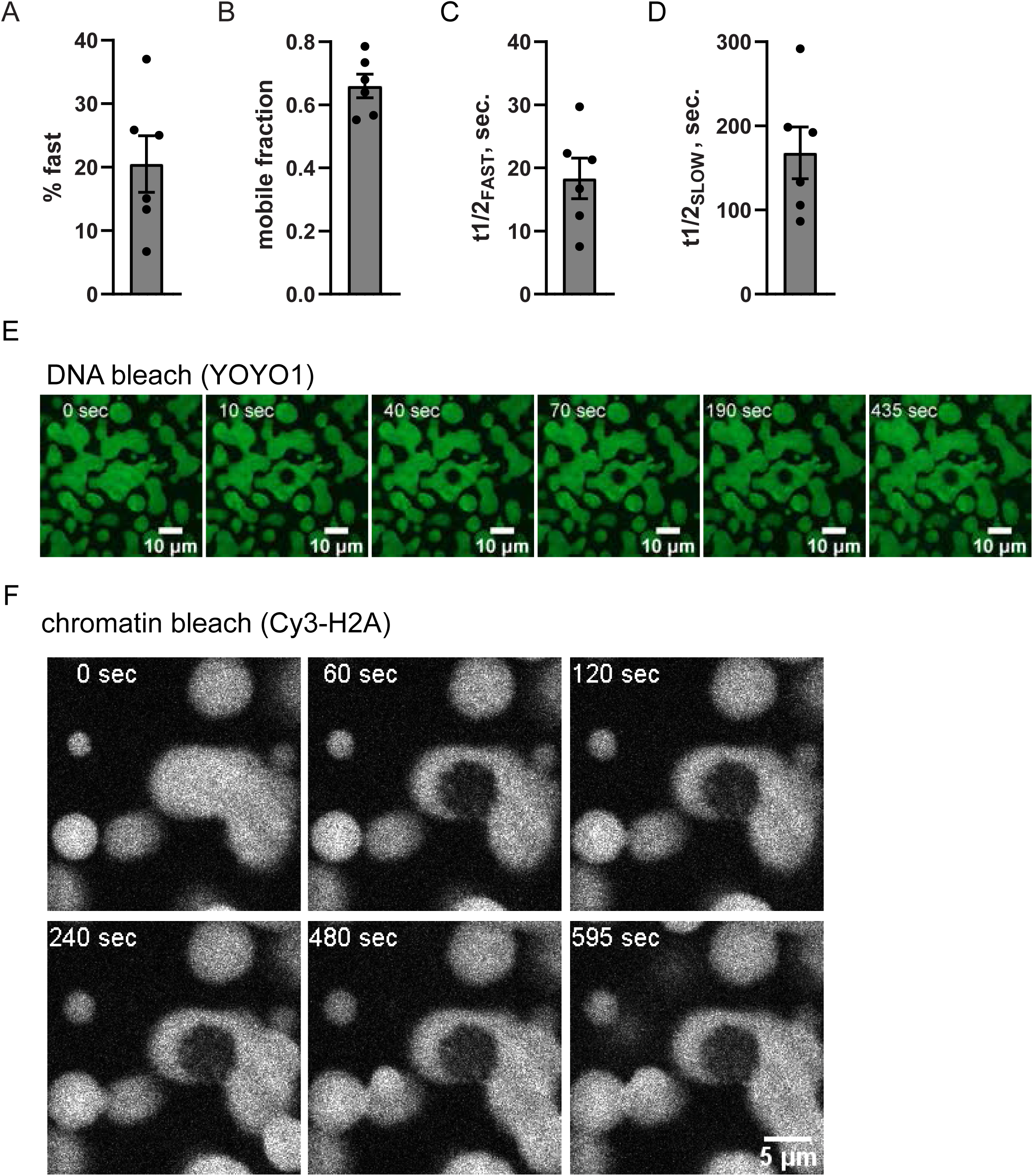
Kinetics of chromatin and Mini-Ph in condensates are distinct. A-D. Summary of parameters from double exponential fits of FRAP data. All bars show mean +/- SEM. E. Time lapse of recovery of chromatin in a condensate. The DNA component of chromatin was visualized (with YOYO1) and bleached. F. Time laps of recovery of chromatin in a condensate after photobleaching. H2A-Cy3 was visualized.

**Supplementary Figure 7.**
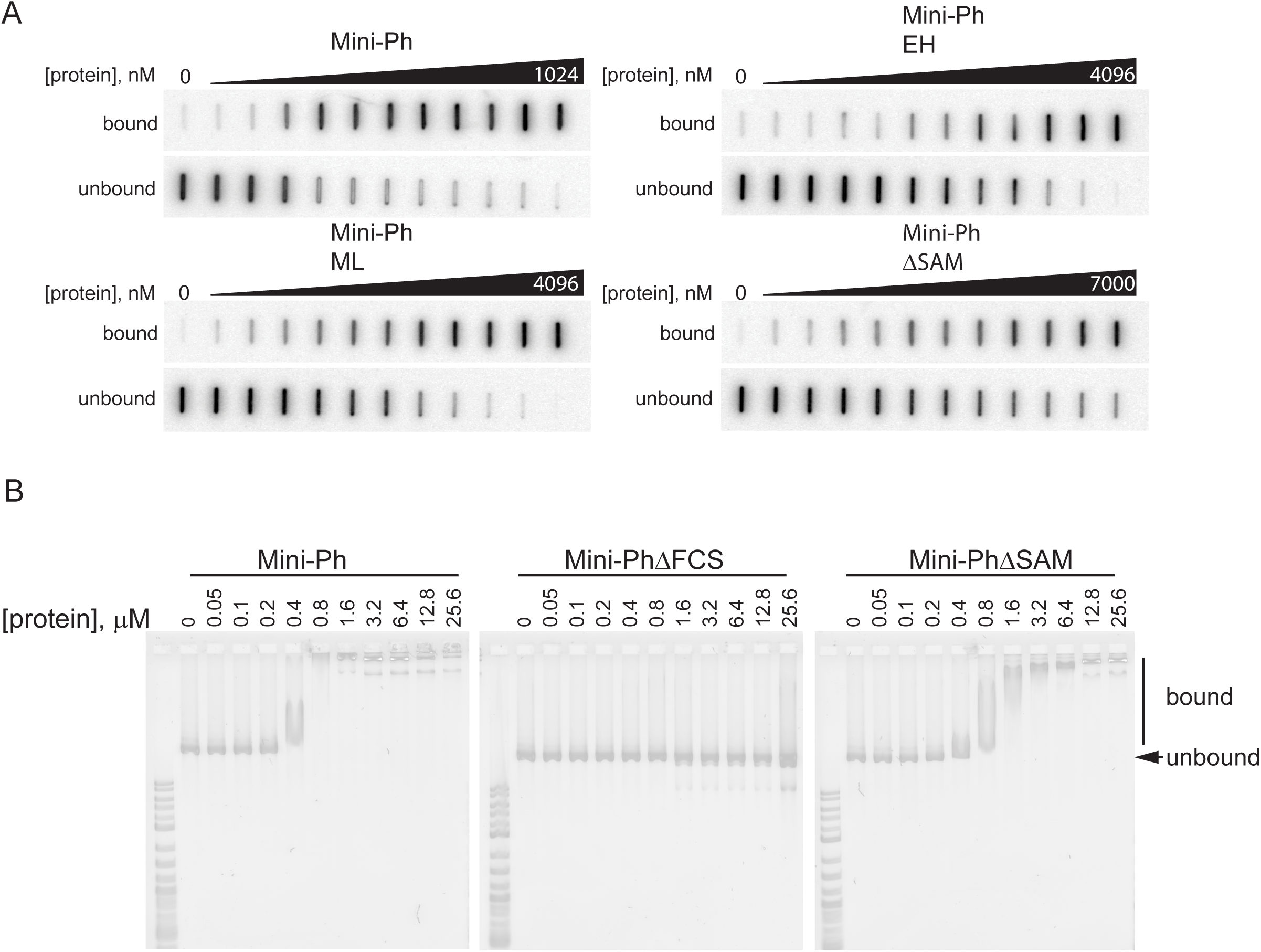
Ph SAM polymerization increases DNA binding affinity of Mini-Ph. A. Representative filters from filter binding experiments to measure DNA binding affinity. B. EMSA demonstrating that Mini-PhΔFCS does not bind DNA. Plasmid DNA was used for EMSA; using these large substrates at high concentrations (25 ng plasmid per reaction) underestimates Mini-Ph DNA binding affinity.

**Supplementary Figure 8.**
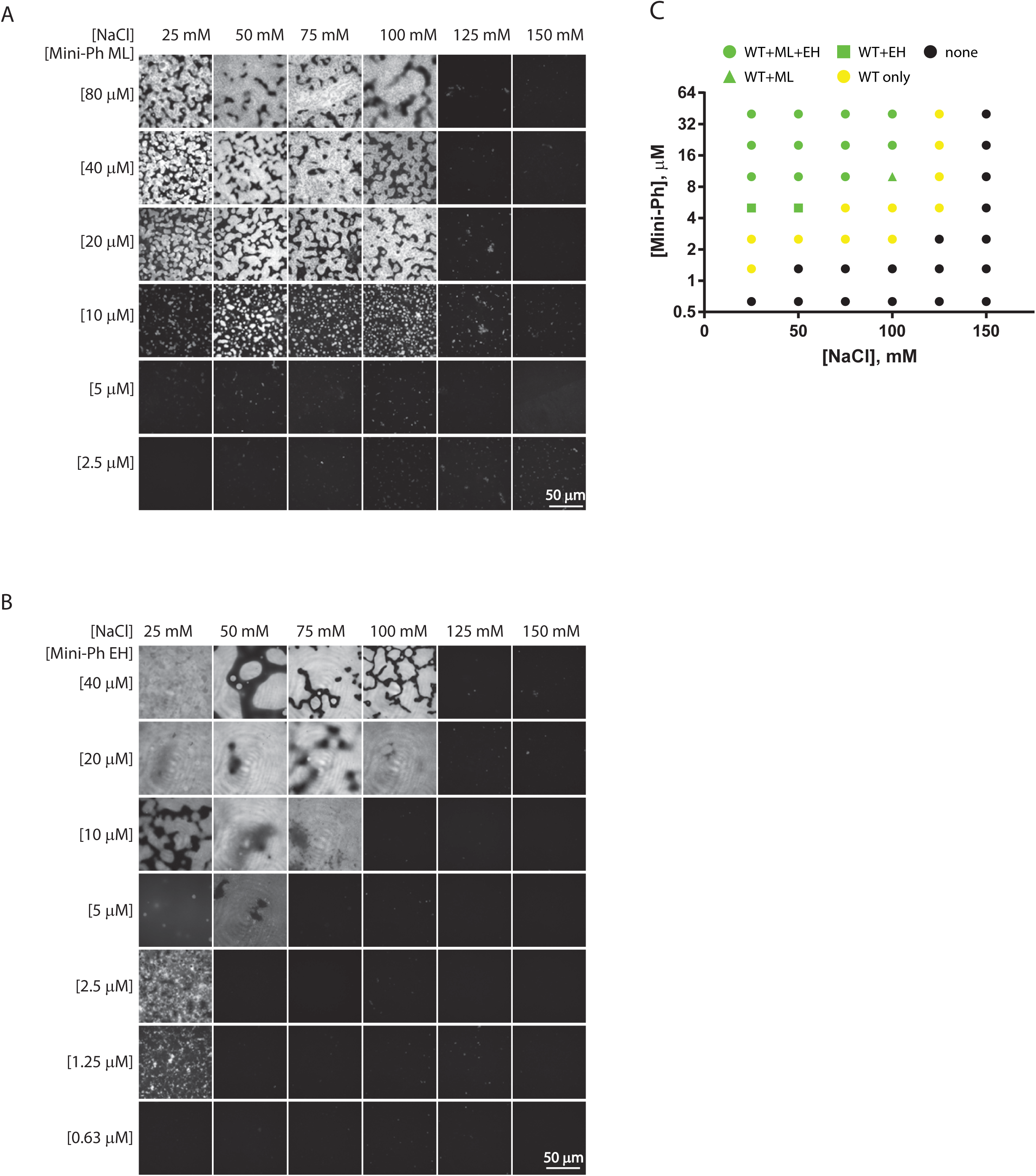
Mini-Ph SAM polymerization mutants are more sensitive to NaCl than wild-type. A, B. Matrix of of Mini-Ph ML (A), and Mini-Ph EH (B)-DNA condensates and across different concentrations of NaCl. C. Plot of one-phase versus two-phase regimens for Mini-Ph, Mini-Ph ML, and Mini-Ph EH. The difference between green and yellow symbols indicates that condensate formation is more sensitive to NaCl for both mutants. See also Supplementary Figure 3 for the matrix of Mini-Ph versus NaCl.

**Supplementary Figure 9.**
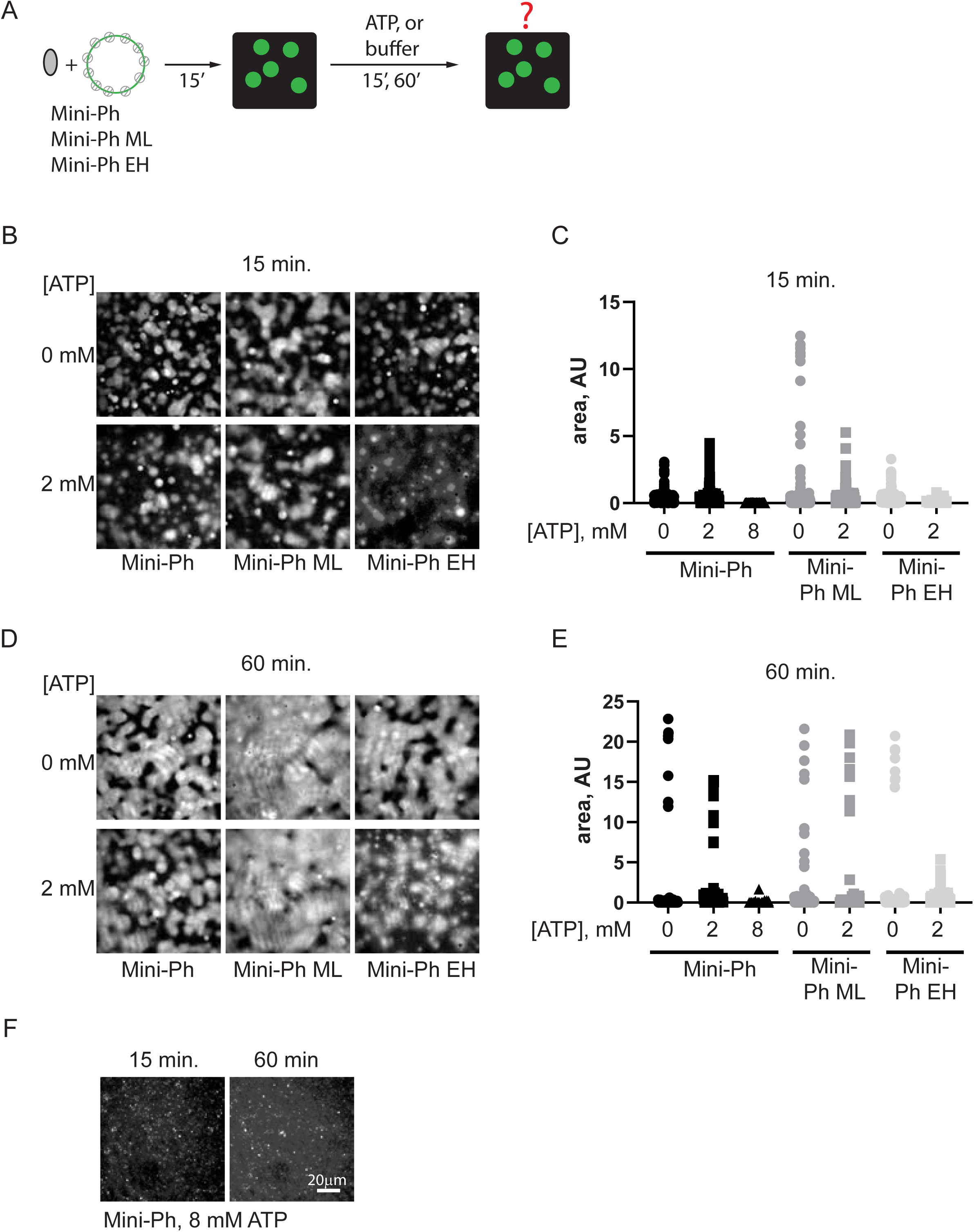
Mini-Ph SAM polymerization mutants are more sensitive to ATP than wild-type Mini-Ph. A. Schematic of experiments to test effect of adding ATP (or buffer as a control) to condensates formed by Mini-Ph, Mini-Ph ML, or Mini-Ph EH with chromatin. B-E. Representative images (B, D) and quantification (C, E) of condensates after 15 min. incubation with ATP or buffer. F. Effect of 8 mM ATP on condensates formed with Mini-Ph and chromatin.

**Supplementary Figure 10.**
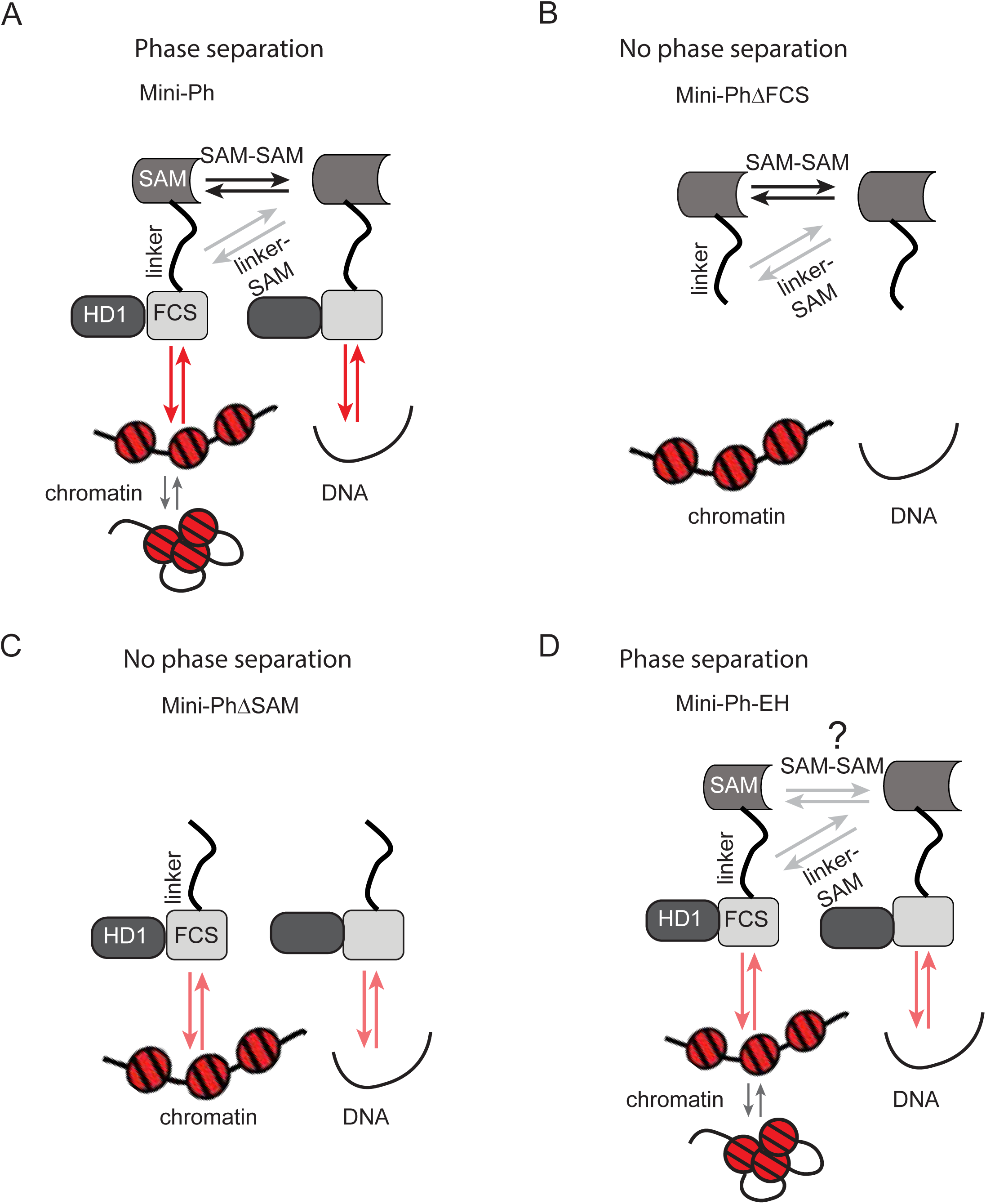
Models for the interactions of Ph SAM in condensate formation. A. Schematic of all known interactions that may occur in Mini-Ph-chromatin or Mini-Ph DNA condensates, including interactions among nucleosomes. B-D. Schematic indicating which interactions are missing when different mutated or truncated proteins that do or do not form condensates are used. The FCS-DNA/chromatin interaction is weaker in the absence of the SAM or when the polymerization interface is disrupted. The residual SAM-SAM interaction in the EH mutant is hypothesized based on the acetylation footprinting results (Supplementary Fig. 13F). The ML mutation, which weakens but does not eliminate SAM-SAM interactions may behave similar to the schematic in A (i.e. the SAM-SAM interaction is dynamic under phase separation conditions, unlike wild-type Mini-Ph which forms short, limited polymers before phase separation). It is important to point out that this is the simplest scheme; it is possible that there are other interactions in the system that have not been characterized. These could include interactions involving the HD1, hinted at by the difference in accessibility measured for Mini-Ph and Mini-Ph EH (Fig. 4D, E). We also do not know how the structure of Mini-Ph polymers influences binding to large chromatin or DNA templates, and whether Mini-Ph binding influences nucleosome-nucleosome interactions (as suggested in the diagrams).

**Supplementary Figure 11.**
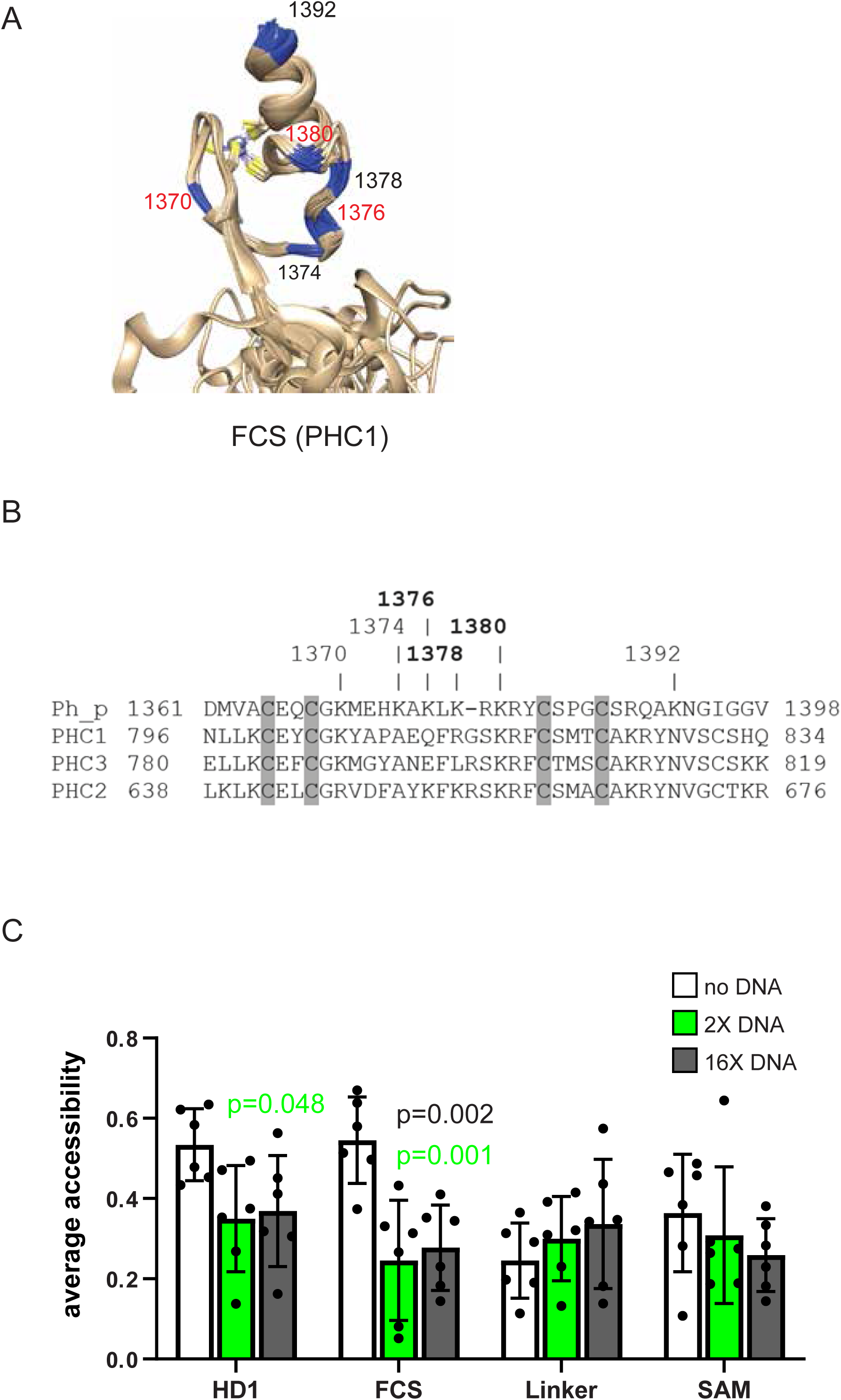
Chemical footprinting analysis of lysine accessibility of Mini-Ph alone and with DNA. A. Structure of the FCS of human PHC1 (PDB 2L8E), with the positions of the equivalent lysine residues in Ph-p indicated. Red numbers indicate positions where accessibility is significantly changed. B. Clustal alignment of human PHCs with Ph-p indicating the relationship between lysines in Ph-p and the sequence of the FCS. C. Average accessibility of lysines in each Mini-Ph region compared across conditions. Accessibility of all residues in each region was averaged for each replicate and the averages compared across conditions by student’t t-test with Holm-Sidak correction for multiple comparisons. Bars show the error +/- SEM. n=6.

**Supplementary Figure 12.**
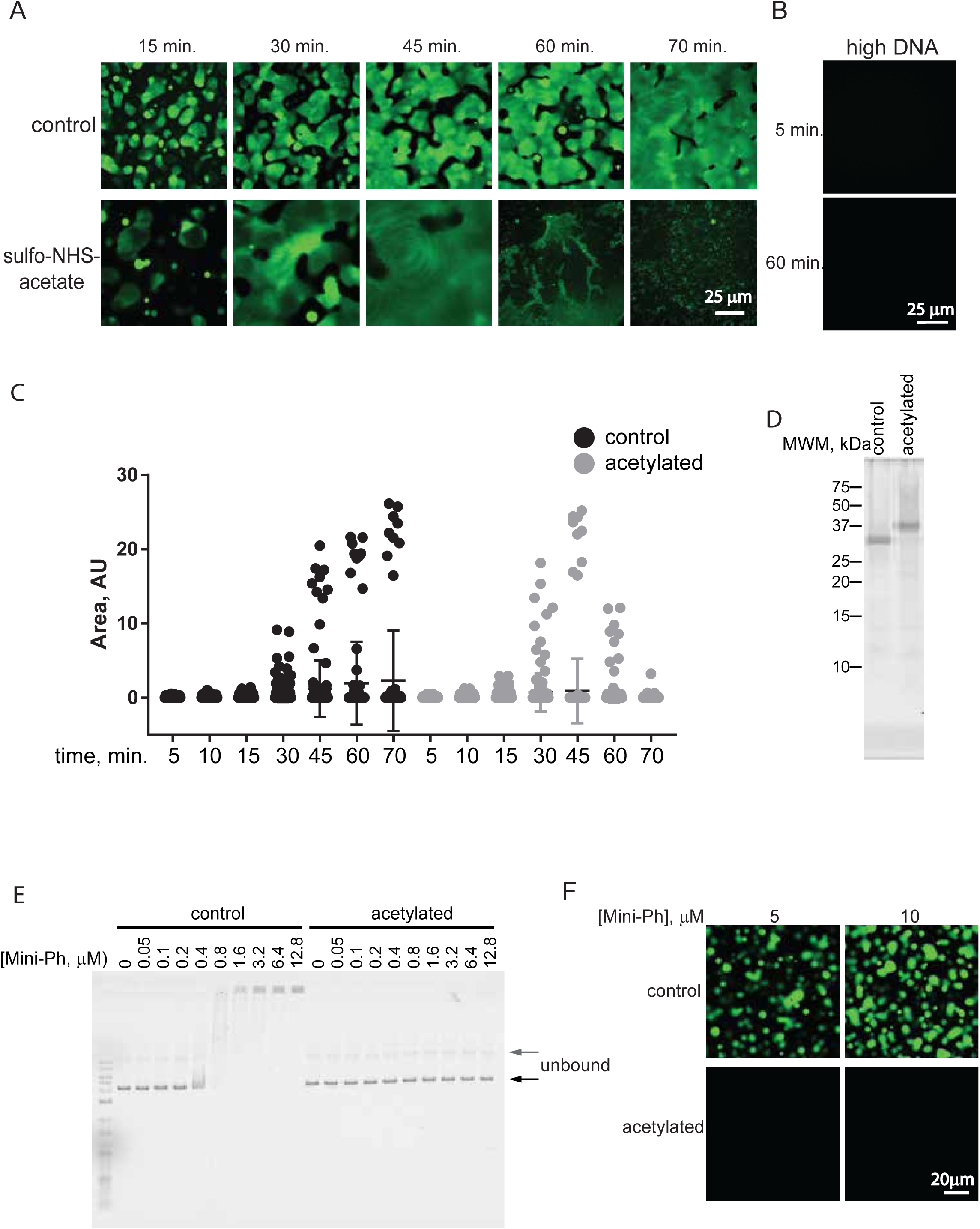
Acetylation of Mini-Ph-DNA blocks condensate formation and DNA binding. Time course of effect of sulfo-NHS acetate on Mini-Ph-DNA condensates. Mass spectrometry samples were collected after 15 min. B. Images of high DNA reaction (without acetylation) in which condensates do not form even after prolonged incubation. C. Quantification of time course shown in A. A similar time course was observed in three experiments. D. SYPRO Ruby stained SDS-PAGE of control Mini-Ph and Mini-Ph after acetylation with sulfo-NHS-Acetate. E. EMSA comparing binding of control and acetylated Mini-Ph to DNA; representative of two experiments. Main unbound band is supercoiled plasmid, and faint band (grey arrow) is nicked. F. Acetylated Mini-Ph does not form condensates with chromatin (representative of three experiments). Images were taken after 60 min. incubation at room temperature.

**Supplementary Figure 13.**
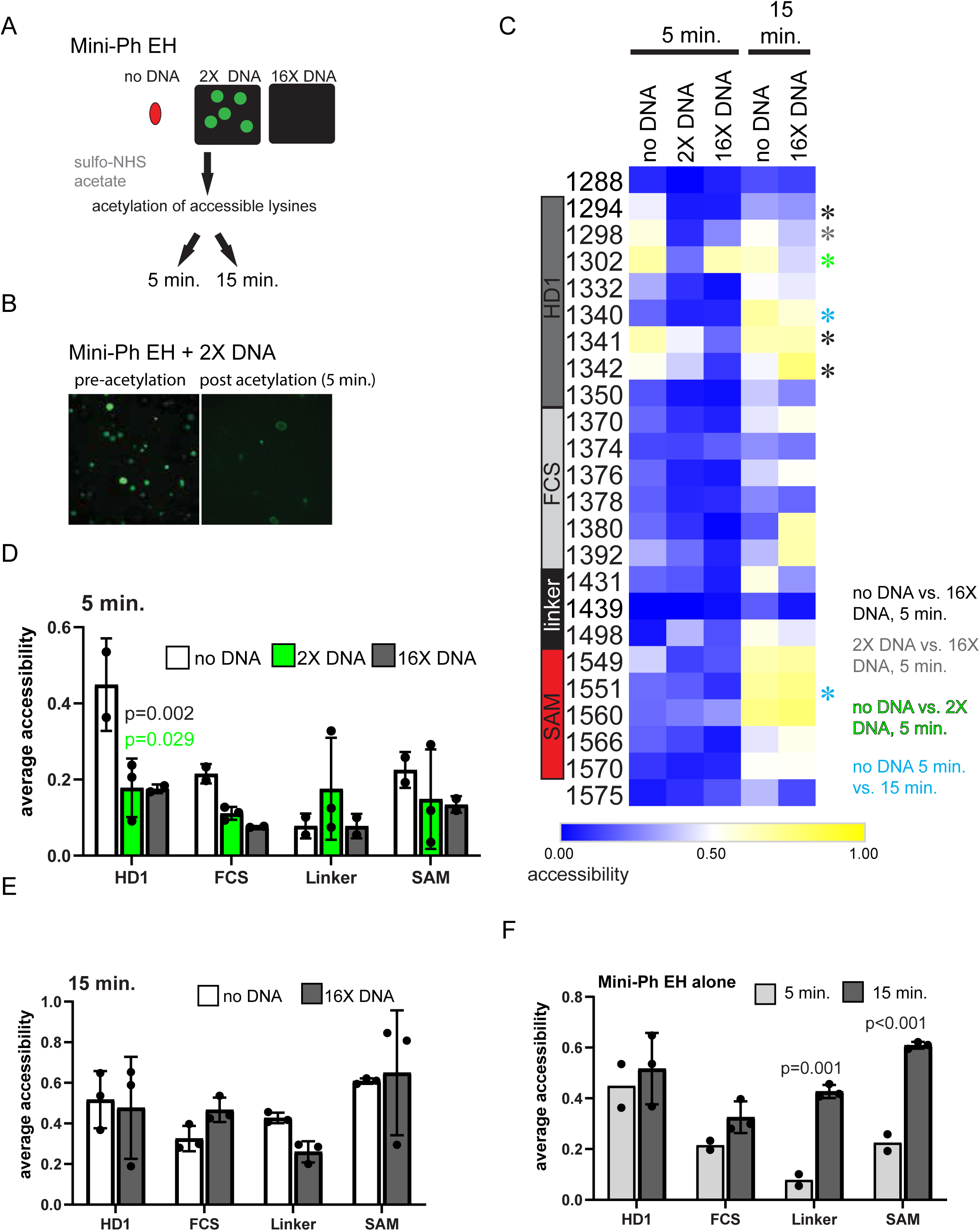
Chemical footprinting analysis of lysine accessibility of polymerization mutant Mini-Ph EH alone and with DNA. A. Schematic of chemical acetylation (abbreviated from Figure 4A). Acetylation was carried out for 5 or 15 min. with Mini-Ph EH because protein-DNA condensates dissolve rapidly after addition of sulfo-NHS acetate. B. Mini-Ph EH-DNA condensates before (left) and after 5 min. of acetylation (right). C. Heat maps of Mini-Ph EH alone, with 2X DNA (condensates form) or 16X DNA (no condensates). Acetylation was carried out for 5 (n=3) or 15 (n=2) min. Asterisks indicate lysines with significantly different accessibility in the indicated comparisons. D-F. Accessibility averaged over Mini-Ph regions for different conditions after 5 min. (D), 15 min. (E), or 5 versus 15 min. for Mini-Ph EH alone (F). Accessibility of all residues in each region was averaged for each replicate and the averages compared across conditions by student’t t-test with Holm-Sidak correction for multiple comparisons. Bars show the error +/- SEM.

**Supplementary Figure 14.**
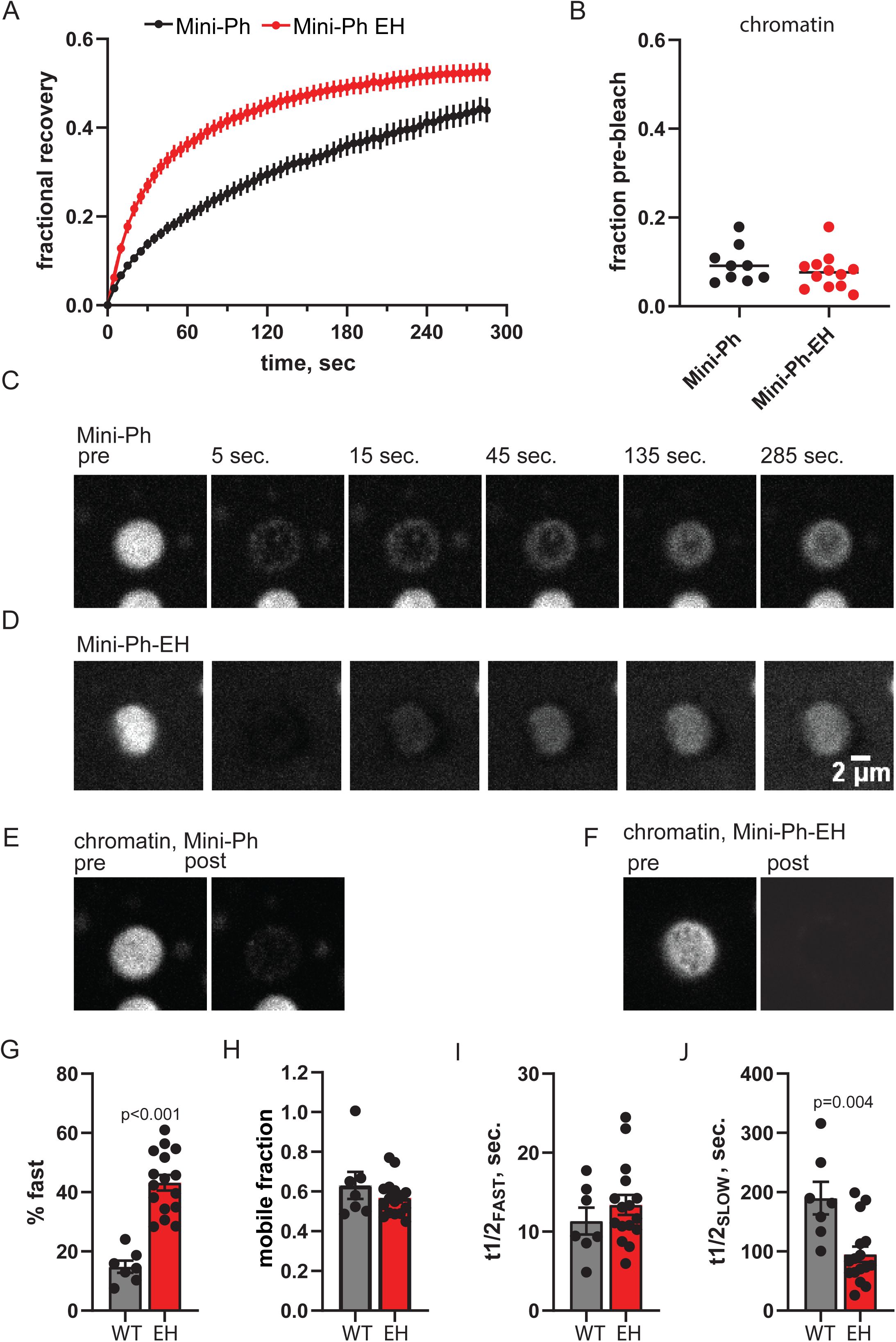
Mini-Ph EH that lacks polymerization activity is more mobile in condensates than wild-type Mini-Ph. A. FRAP traces for Mini-Ph and Mini-Ph EH with chromatin. Representative traces with R^2^ for fits >0.99 were pooled for condensates formed at different times (22-41 min. for Mini-Ph; 28-73 min. for Mini-Ph-EH; n=5). Graph shows the mean +/- SEM. Data were fit with a double exponential equation. B. Fluorescence of chromatin (H2A-Cy3) before and after FRAP experiments. The same regions of interest (ROI) used to measure FRAP, acquisition induced bleaching, and background were used to analyze chromatin as for Mini-Ph. We were unable to collect FRAP traces in both channels simultaneously. C, D. Representative bleaching and recovery of single condensate for Alexa-647 labelled Mini-Ph (C) and Mini-Ph-EH. Condensates were formed for 25 min. for Mini-Ph and 28 min. for Mini-Ph-EH. E, F. Fluorescence of chromatin (H2A-Cy3) for the same condensates shown in C and D immediately before and after the FRAP experiment. G-I Summary of parameters from double exponential fits. All graphs show the mean +/-SEM. Mini-Ph and Mini-PH EH were compared by Mann-Whitney test. All fits have R^2^>0.99.

**Supplementary Figure 15.**
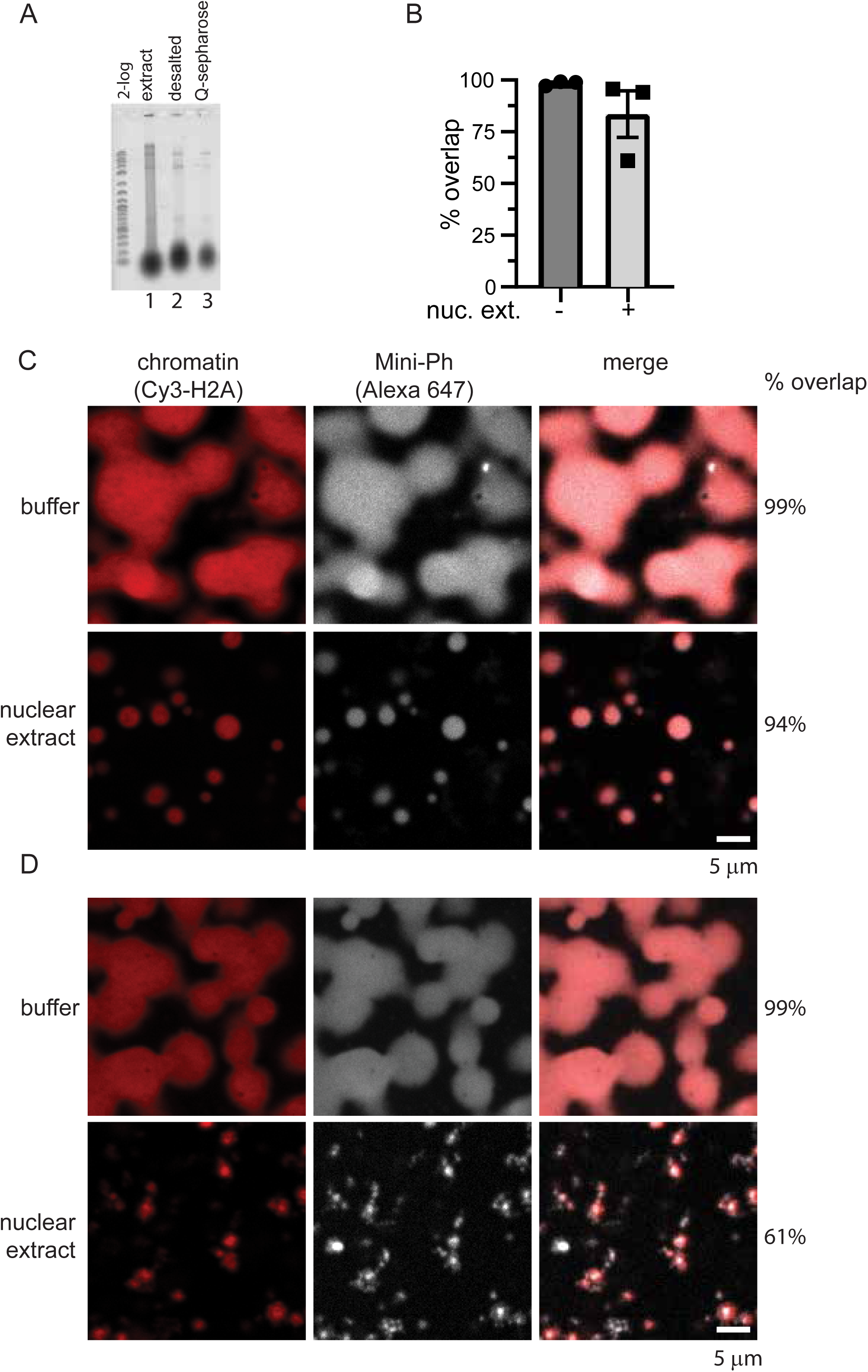
Colocalization analysis of Mini-Ph and chromatin in nuclear extracts. A. SYBR Gold stained gel of nuclear extract (lane 1), extract after desalting (lane 2), and after incubation with Q-sepharose to deplete nucleic acids (lane 3). B. Summary of overlap between Mini-Ph condensates and chromatin incubated in buffer or in nuclear extract from three experiments. C, D. Representative images from two of the three experiments; two resembled panel C, while in one experiment, Mini-Ph structures that are not positive for Cy3-H2A are observed. These structures frequently appear to be connected to chromatin condensates.

**Supplementary Figure 16.**
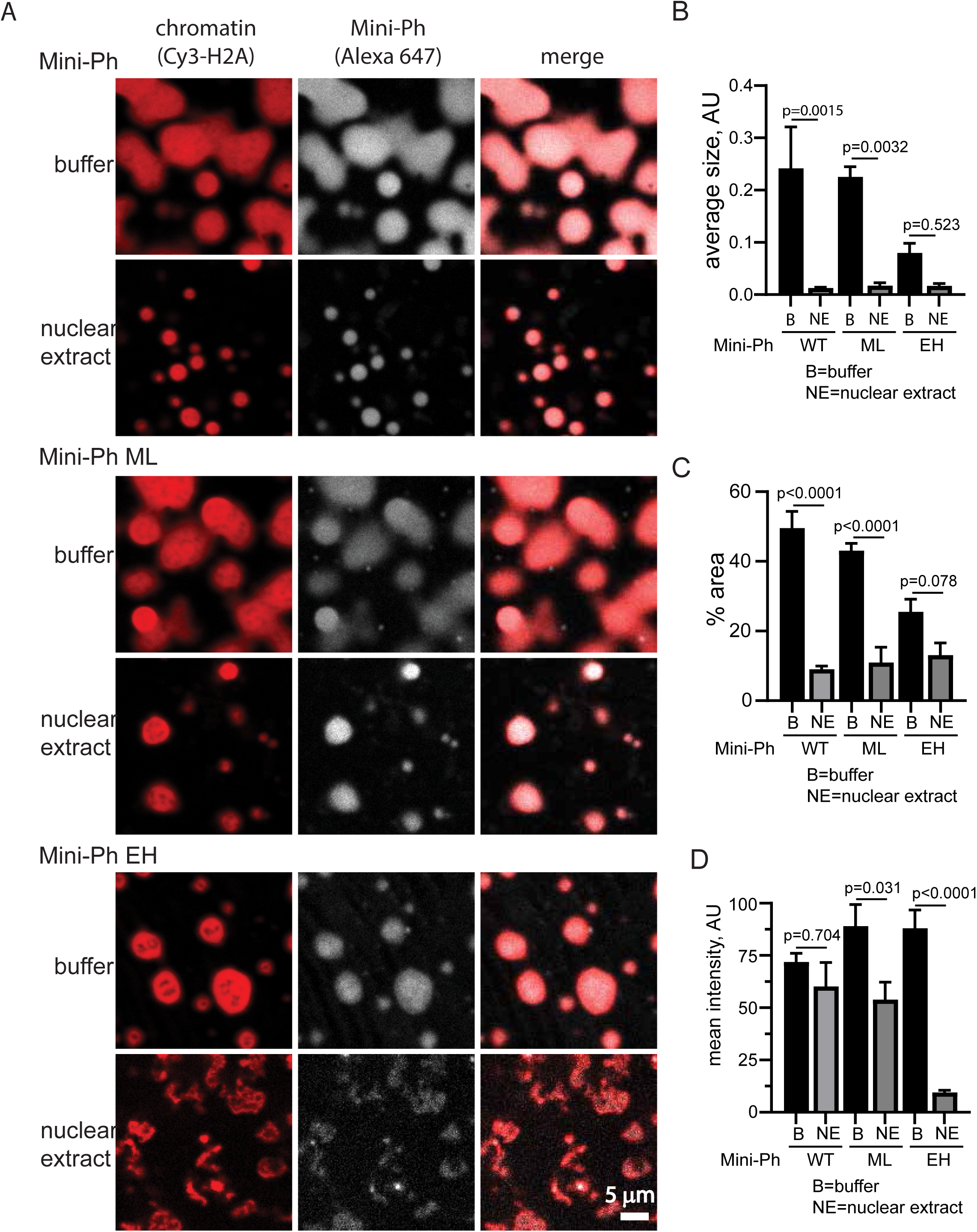
Effect of nuclear extracts on condensates formed by Mini-Ph, or Mini-Ph polymerization mutants with chromatin. A. Representative images of condensates incubated in buffer (top panel in each set), or nuclear extracts (bottom panel in each set). Image intensities were adjusted to facilitate visualization of the structures. The structures formed by Mini-Ph EH with chromatin in nuclear extract are much fainter than those formed in buffer are. B-D. Average size of structures (B), % area covered by condensates (C), or mean intensity of structures (D) after incubation in buffer or extract from three experiments. Bars are the mean +SEM and p-values are for ANOVA comparing buffer to nuclear extract for each protein with Sidak correction for multiple comparisons.

**Supplementary Figure 17.**
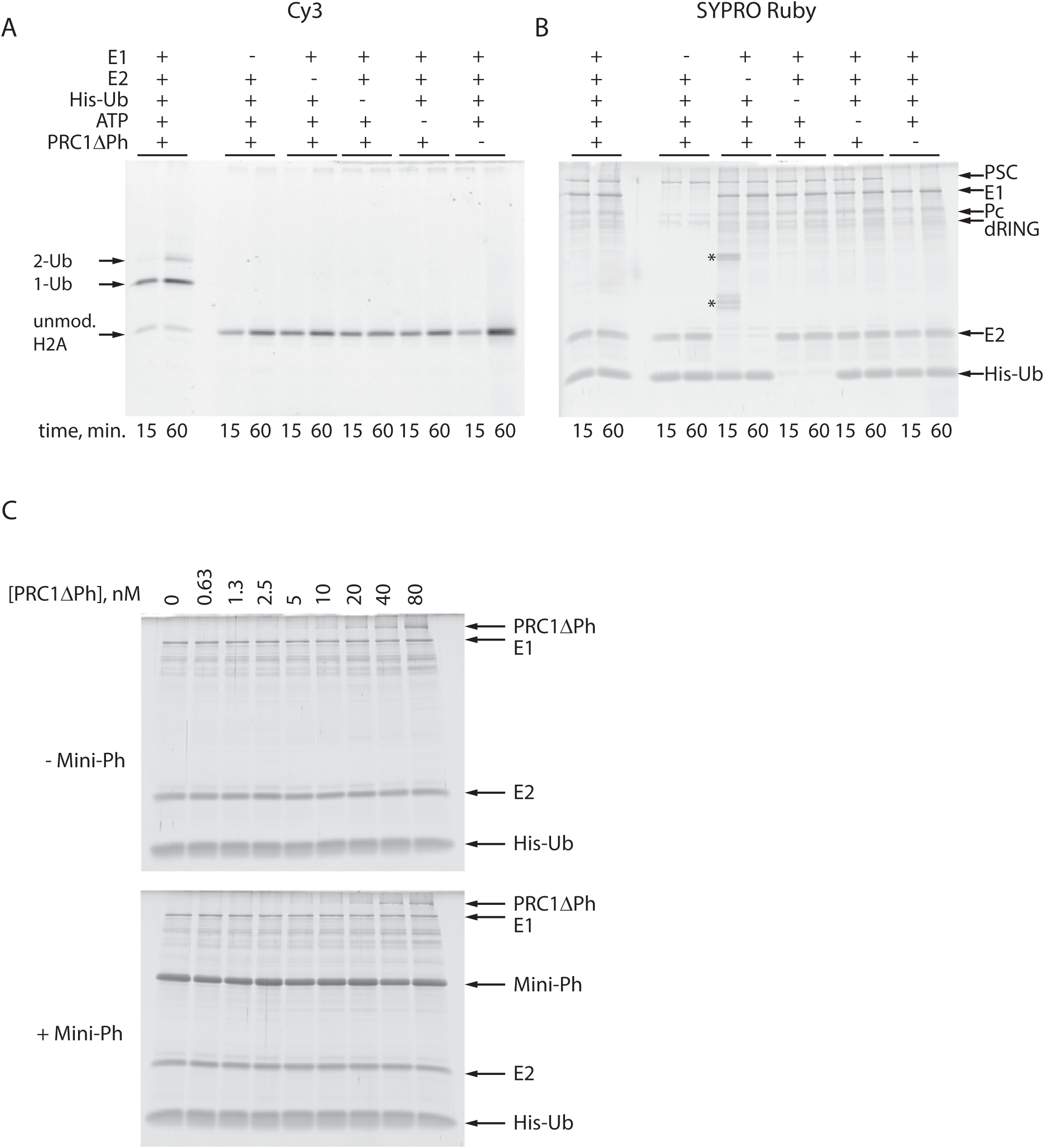
Histone ubiquitylation assay. A, B. SDS-PAGE of histone ubiquitylation assay. A shows Cy3-labelled histone H2A and B is same gel after SYPRO-Ruby staining. The E1 and E2, ATP, Ubiquitin, and PRC1ΔPh are all required for ubiquitylation. *contaminants present in one gel lane. C. SYPRO Ruby stain of gels shown in Figure 6.

**Supplementary Figure 18.**
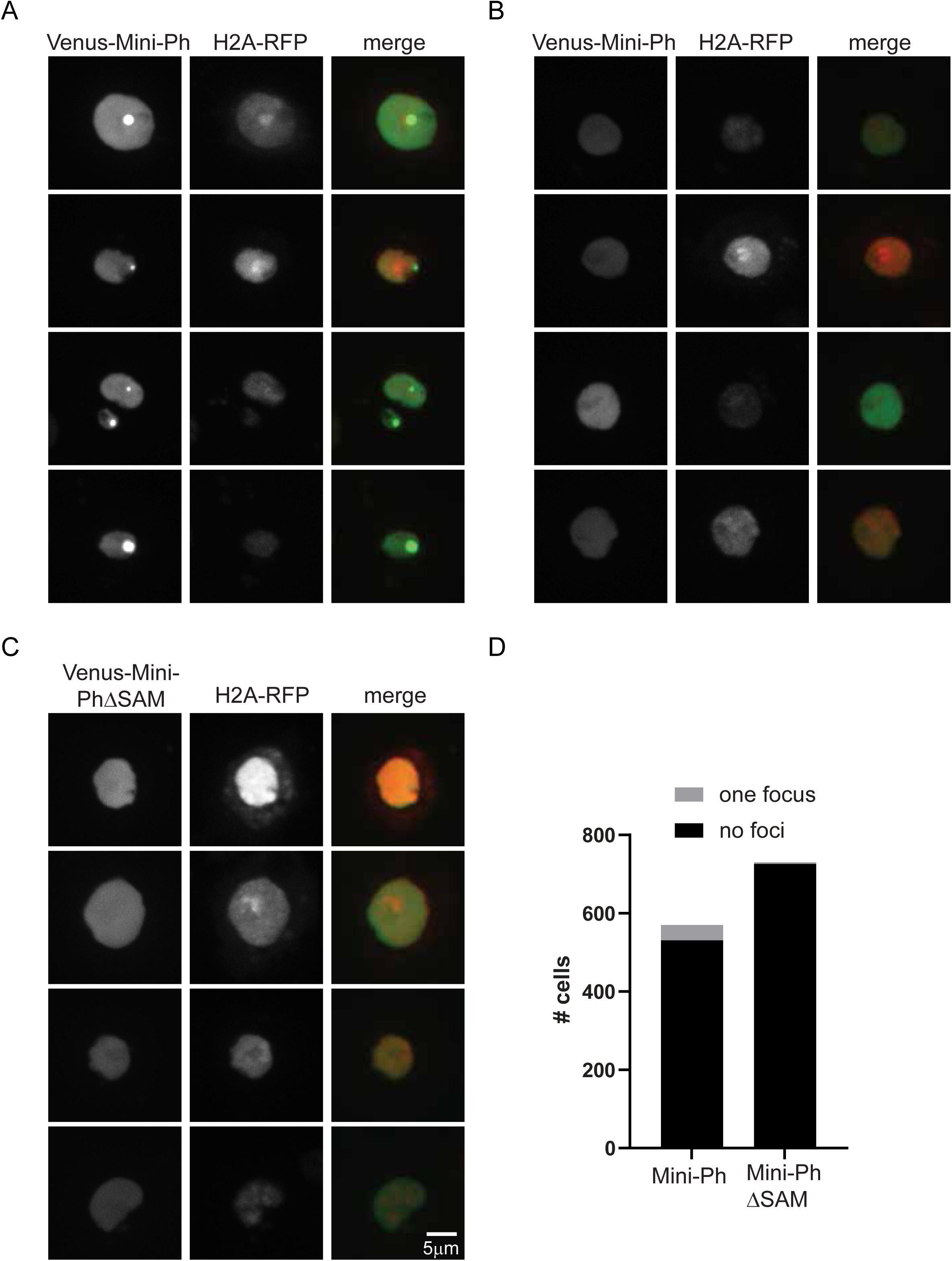
Mini-Ph forms a single, SAM-dependent focus or no foci in cells. A-C. Representative images of S2R+ cells transfected with Venus-Mini-Ph (A, B) or Venus-Mini-PhΔSAM (C) and H2A-RFP. Cells expressing Venus-Mini-Ph either have a single focus, as in A, or no foci, as in B. We do not observe multiple Mini-Ph foci in cells. D. Quantification of cells with foci. 7% of Venus-Mini-Ph (39/570) and 0.5% of Venus-Mini-PhΔSAM (4/730) cells were identified as having a single maxima (focus). In the case of Mini-PhDSAM, the small number of identified foci represent cells with very high expression rather than actual foci.

## Notes

### Competing Interest Statement

The authors have declared no competing interest.

## References

1 Di Croce, L. & Helin, K. Transcriptional regulation by Polycomb group proteins. Nat Struct Mol Biol 20, 1147–1155, doi:10.1038/nsmb.2669 (2013).

2 Entrevan, M., Schuettengruber, B. & Cavalli, G. Regulation of Genome Architecture and Function by Polycomb Proteins. Trends Cell Biol 26, 511–525, doi:10.1016/j.tcb.2016.04.009 (2016).

3 Kassis, J. A., Kennison, J. A. & Tamkun, J. W. Polycomb and Trithorax Group Genes in Drosophila. Genetics 206, 1699–1725, doi:10.1534/genetics.115.185116 (2017).

4 Loubiere, V., Martinez, A. M. & Cavalli, G. Cell Fate and Developmental Regulation Dynamics by Polycomb Proteins and 3D Genome Architecture. Bioessays 41, e1800222, doi:10.1002/bies.201800222 (2019).

5 Robinson, A. K. et al. The growth-suppressive function of the polycomb group protein polyhomeotic is mediated by polymerization of its sterile alpha motif (SAM) domain. J Biol Chem 287, 8702–8713, doi:10.1074/jbc.M111.336115 (2012).

6 Kang, J. J., Faubert, D., Boulais, J. & Francis, N. J. DNA Binding Reorganizes the Intrinsically Disordered C-Terminal Region of PSC in Drosophila PRC1. J Mol Biol, doi:10.1016/j.jmb.2020.07.002 (2020).

7 Fursova, N. A. et al. Synergy between Variant PRC1 Complexes Defines Polycomb-Mediated Gene Repression. Mol Cell 74, 1020–1036 e1028, doi:10.1016/j.molcel.2019.03.024 (2019).

8 Scelfo, A. et al. Functional Landscape of PCGF Proteins Reveals Both RING1A/B-Dependent-and RING1A/B-Independent-Specific Activities. Mol Cell 74, 1037–1052 e1037, doi:10.1016/j.molcel.2019.04.002 (2019).

9 Gao, Z. et al. PCGF homologs, CBX proteins, and RYBP define functionally distinct PRC1 family complexes. Mol Cell 45, 344–356, doi:10.1016/j.molcel.2012.01.002 (2012).

10 Pemberton, H. et al. Genome-wide co-localization of Polycomb orthologs and their effects on gene expression in human fibroblasts. Genome Biol 15, R23, doi:10.1186/gb-2014-15-2-r23 (2014).

11 Francis, N. J., Kingston, R. E. & Woodcock, C. L. Chromatin compaction by a polycomb group protein complex. Science 306, 1574–1577, doi:10.1126/science.1100576 (2004).

12 Cheutin, T. & Cavalli, G. Loss of PRC1 induces higher-order opening of Hox loci independently of transcription during Drosophila embryogenesis. Nat Commun 9, 3898, doi:10.1038/s41467-018-05945-4 (2018).

13 Isono, K. et al. SAM domain polymerization links subnuclear clustering of PRC1 to gene silencing. Dev Cell 26, 565–577, doi:10.1016/j.devcel.2013.08.016 (2013).

14 Wani, A. H. et al. Chromatin topology is coupled to Polycomb group protein subnuclear organization. Nat Commun 7, 10291, doi:10.1038/ncomms10291 (2016).

15 Boettiger, A. N. et al. Super-resolution imaging reveals distinct chromatin folding for different epigenetic states. Nature 529, 418–422, doi:10.1038/nature16496 (2016).

16 Eskeland, R. et al. Ring1B compacts chromatin structure and represses gene expression independent of histone ubiquitination. Mol Cell 38, 452–464, doi:10.1016/j.molcel.2010.02.032 (2010).

17 Boyle, S. et al. A central role for canonical PRC1 in shaping the 3D nuclear landscape. Genes Dev 34, 931–949, doi:10.1101/gad.336487.120 (2020).

18 Kim, C. A., Gingery, M., Pilpa, R. M. & Bowie, J. U. The SAM domain of polyhomeotic forms a helical polymer. Nat Struct Biol 9, 453–457, doi:10.1038/nsb802 (2002).

19 Kim, C. A. & Bowie, J. U. SAM domains: uniform structure, diversity of function. Trends Biochem Sci 28, 625–628, doi:10.1016/j.tibs.2003.11.001 (2003).

20 Gambetta, M. C. & Muller, J. O-GlcNAcylation prevents aggregation of the Polycomb group repressor polyhomeotic. Dev Cell 31, 629–639, doi:10.1016/j.devcel.2014.10.020 (2014).

21 Tsuboi, M. et al. Ubiquitination-Independent Repression of PRC1 Targets during Neuronal Fate Restriction in the Developing Mouse Neocortex. Dev Cell 47, 758–772 e755, doi:10.1016/j.devcel.2018.11.018 (2018).

22 Cheutin, T. & Cavalli, G. Progressive polycomb assembly on H3K27me3 compartments generates polycomb bodies with developmentally regulated motion. PLoS Genet 8, e1002465, doi:10.1371/journal.pgen.1002465 (2012).

23 Banani, S. F., Lee, H. O., Hyman, A. A. & Rosen, M. K. Biomolecular condensates: organizers of cellular biochemistry. Nat Rev Mol Cell Biol 18, 285–298, doi:10.1038/nrm.2017.7 (2017).

24 Boeynaems, S. et al. Protein Phase Separation: A New Phase in Cell Biology. Trends Cell Biol 28, 420–435, doi:10.1016/j.tcb.2018.02.004 (2018).

25 Shin, Y. & Brangwynne, C. P. Liquid phase condensation in cell physiology and disease. Science 357, doi:10.1126/science.aaf4382 (2017).

26 Strom, A. R. & Brangwynne, C. P. The liquid nucleome - phase transitions in the nucleus at a glance. J Cell Sci 132, doi:10.1242/jcs.235093 (2019).

27 Fay, M. M. & Anderson, P. J. The Role of RNA in Biological Phase Separations. J Mol Biol 430, 4685–4701, doi:10.1016/j.jmb.2018.05.003 (2018).

28 Gibson, B. A. et al. Organization of Chromatin by Intrinsic and Regulated Phase Separation. Cell 179, 470–484 e421, doi:10.1016/j.cell.2019.08.037 (2019).

29 Huo, X. et al. The Nuclear Matrix Protein SAFB Cooperates with Major Satellite RNAs to Stabilize Heterochromatin Architecture Partially through Phase Separation. Mol Cell 77, 368–383 e367, doi:10.1016/j.molcel.2019.10.001 (2020).

30 Larson, A. G. et al. Liquid droplet formation by HP1alpha suggests a role for phase separation in heterochromatin. Nature 547, 236–240, doi:10.1038/nature22822 (2017).

31 Maeshima, K. et al. Nucleosomal arrays self-assemble into supramolecular globular structures lacking 30-nm fibers. EMBO J 35, 1115–1132, doi:10.15252/embj.201592660 (2016).

32 Strom, A. R. et al. Phase separation drives heterochromatin domain formation. Nature 547, 241–245, doi:10.1038/nature22989 (2017).

33 Wang, L. et al. Histone Modifications Regulate Chromatin Compartmentalization by Contributing to a Phase Separation Mechanism. Mol Cell 76, 646–659 e646, doi:10.1016/j.molcel.2019.08.019 (2019).

34 Hnisz, D., Shrinivas, K., Young, R. A., Chakraborty, A. K. & Sharp, P. A. A Phase Separation Model for Transcriptional Control. Cell 169, 13–23, doi:10.1016/j.cell.2017.02.007 (2017).

35 McSwiggen, D. T., Mir, M., Darzacq, X. & Tjian, R. Evaluating phase separation in live cells: diagnosis, caveats, and functional consequences. Genes Dev 33, 1619–1634, doi:10.1101/gad.331520.119 (2019).

36 Boija, A. et al. Transcription Factors Activate Genes through the Phase-Separation Capacity of Their Activation Domains. Cell 175, 1842–1855 e1816, doi:10.1016/j.cell.2018.10.042 (2018).

37 Sabari, B. R. et al. Coactivator condensation at super-enhancers links phase separation and gene control. Science 361, doi:10.1126/science.aar3958 (2018).

38 Kilic, S. et al. Phase separation of 53BP1 determines liquid-like behavior of DNA repair compartments. EMBO J 38, e101379, doi:10.15252/embj.2018101379 (2019).

39 Oshidari, R. et al. DNA repair by Rad52 liquid droplets. Nat Commun 11, 695, doi:10.1038/s41467-020-14546-z (2020).

40 Plys, A. J. et al. Phase separation of Polycomb-repressive complex 1 is governed by a charged disordered region of CBX2. Genes Dev 33, 799–813, doi:10.1101/gad.326488.119 (2019).

41 Qu, W., Wang, Z. & Zhang, H. Phase separation of the C. elegans Polycomb protein SOP-2 is modulated by RNA and sumoylation. Protein Cell, doi:10.1007/s13238-019-00680-y (2020).

42 Tatavosian, R. et al. Nuclear condensates of the Polycomb protein chromobox 2 (CBX2) assemble through phase separation. J Biol Chem 294, 1451–1463, doi:10.1074/jbc.RA118.006620 (2019).

43 Shin, Y. et al. Liquid Nuclear Condensates Mechanically Sense and Restructure the Genome. Cell 175, 1481–1491 e1413, doi:10.1016/j.cell.2018.10.057 (2018).

44 Erdel, F. et al. Mouse Heterochromatin Adopts Digital Compaction States without Showing Hallmarks of HP1-Driven Liquid-Liquid Phase Separation. Mol Cell 78, 236–249 e237, doi:10.1016/j.molcel.2020.02.005 (2020).

45 Hodgson, J. W. et al. The polyhomeotic locus of Drosophila melanogaster is transcriptionally and post-transcriptionally regulated during embryogenesis. Mech Dev 66, 69–81, doi:10.1016/s0925-4773(97)00091-9 (1997).

46 Wang, R. et al. Identification of nucleic acid binding residues in the FCS domain of the polycomb group protein polyhomeotic. Biochemistry 50, 4998–5007, doi:10.1021/bi101487s (2011).

47 Simpson, R. T., Thoma, F. & Brubaker, J. M. Chromatin reconstituted from tandemly repeated cloned DNA fragments and core histones: a model system for study of higher order structure. Cell 42, 799–808, doi:10.1016/0092-8674(85)90276-4 (1985).

48 Harmon, T. S., Holehouse, A. S., Rosen, M. K. & Pappu, R. V. Intrinsically disordered linkers determine the interplay between phase separation and gelation in multivalent proteins. Elife 6, doi:10.7554/eLife.30294 (2017).

49 Choi, J. M., Holehouse, A. S. & Pappu, R. V. Physical Principles Underlying the Complex Biology of Intracellular Phase Transitions. Annu Rev Biophys 49, 107–133, doi:10.1146/annurev-biophys-121219-081629 (2020).

50 Robinson, A. K. et al. Human polyhomeotic homolog 3 (PHC3) sterile alpha motif (SAM) linker allows open-ended polymerization of PHC3 SAM. Biochemistry 51, 5379–5386, doi:10.1021/bi3004318 (2012).

51 Taylor, N. O., Wei, M. T., Stone, H. A. & Brangwynne, C. P. Quantifying Dynamics in Phase-Separated Condensates Using Fluorescence Recovery after Photobleaching. Biophys J 117, 1285–1300, doi:10.1016/j.bpj.2019.08.030 (2019).

52 Fonseca, J. P. et al. In vivo Polycomb kinetics and mitotic chromatin binding distinguish stem cells from differentiated cells. Genes Dev 26, 857–871, doi:10.1101/gad.184648.111 (2012).

53 Boeynaems, S. et al. Spontaneous driving forces give rise to protein-RNA condensates with coexisting phases and complex material properties. Proc Natl Acad Sci U S A 116, 7889–7898, doi:10.1073/pnas.1821038116 (2019).

54 Frank, L. & Rippe, K. Repetitive RNAs as Regulators of Chromatin-Associated Subcompartment Formation by Phase Separation. J Mol Biol, doi:10.1016/j.jmb.2020.04.015 (2020).

55 Strickfaden, H. et al. Condensed chromatin behaves like a solid on the mesoscale in vitro and in living cells. bioRxiv, doi: https://doi.org/10.1101/2020.05.06.079905 (2020).

56 Patel, A. et al. ATP as a biological hydrotrope. Science 356, 753–756, doi:10.1126/science.aaf6846 (2017).

57 Bratek-Skicki, A., Pancsa, R., Meszaros, B., Van Lindt, J. & Tompa, P. A guide to regulation of the formation of biomolecular condensates. FEBS J 287, 1924–1935, doi:10.1111/febs.15254 (2020).

58 Alfieri, C. et al. Structural basis for targeting the chromatin repressor Sfmbt to Polycomb response elements. Genes Dev 27, 2367–2379, doi:10.1101/gad.226621.113 (2013).

59 Gambetta, M. C., Oktaba, K. & Muller, J. Essential role of the glycosyltransferase sxc/Ogt in polycomb repression. Science 325, 93–96, doi:10.1126/science.1169727 (2009).

60 Bonnet, J. et al. Quantification of Proteins and Histone Marks in Drosophila Embryos Reveals Stoichiometric Relationships Impacting Chromatin Regulation. Dev Cell 51, 632–644 e636, doi:10.1016/j.devcel.2019.09.011 (2019).

61 Zhang, H. et al. Global regulation of Hox gene expression in C. elegans by a SAM domain protein. Dev Cell 4, 903–915, doi:10.1016/s1534-5807(03)00136-9 (2003).

62 Zhang, H. et al. SUMO modification is required for in vivo Hox gene regulation by the Caenorhabditis elegans Polycomb group protein SOP-2. Nat Genet 36, 507–511, doi:10.1038/ng1336 (2004).

63 Zhang, H. et al. The C. elegans Polycomb gene SOP-2 encodes an RNA binding protein. Mol Cell 14, 841–847, doi:10.1016/j.molcel.2004.06.001 (2004).

64 Erdel, F. & Rippe, K. Formation of Chromatin Subcompartments by Phase Separation. Biophys J 114, 2262–2270, doi:10.1016/j.bpj.2018.03.011 (2018).

65 Hildebrand, E. M. & Dekker, J. Mechanisms and Functions of Chromosome Compartmentalization. Trends Biochem Sci 45, 385–396, doi:10.1016/j.tibs.2020.01.002 (2020).

66 Shrinivas, K. et al. Enhancer Features that Drive Formation of Transcriptional Condensates. Mol Cell 75, 549–561 e547, doi:10.1016/j.molcel.2019.07.009 (2019).

67 Oksuz, O. et al. Capturing the Onset of PRC2-Mediated Repressive Domain Formation. Mol Cell 70, 1149–1162 e1145, doi:10.1016/j.molcel.2018.05.023 (2018).

68 Kim, C. A. et al. Polymerization of the SAM domain of TEL in leukemogenesis and transcriptional repression. EMBO J 20, 4173–4182, doi:10.1093/emboj/20.15.4173 (2001).

69 Francis, N. J., Saurin, A. J., Shao, Z. & Kingston, R. E. Reconstitution of a functional core polycomb repressive complex. Mol Cell 8, 545–556, doi:10.1016/s1097-2765(01)00316-1 (2001).

70 Taherbhoy, A. M., Huang, O. W. & Cochran, A. G. BMI1-RING1B is an autoinhibited RING E3 ubiquitin ligase. Nat Commun 6, 7621, doi:10.1038/ncomms8621 (2015).

71 Zhao, J. et al. RYBP/YAF2-PRC1 complexes and histone H1-dependent chromatin compaction mediate propagation of H2AK119ub1 during cell division. Nat Cell Biol 22, 439–452, doi:10.1038/s41556-020-0484-1 (2020).

72 Gallego, L. D. et al. Phase separation directs ubiquitination of gene-body nucleosomes. Nature 579, 592–597, doi:10.1038/s41586-020-2097-z (2020).

73 Sanulli, S. et al. HP1 reshapes nucleosome core to promote phase separation of heterochromatin. Nature 575, 390–394, doi:10.1038/s41586-019-1669-2 (2019).

74 Gutierrez, L. et al. The role of the histone H2A ubiquitinase Sce in Polycomb repression. Development 139, 117–127, doi:10.1242/dev.074450 (2012).

75 Pengelly, A. R., Kalb, R., Finkl, K. & Muller, J. Transcriptional repression by PRC1 in the absence of H2A monoubiquitylation. Genes Dev 29, 1487–1492, doi:10.1101/gad.265439.115 (2015).

76 Lee, H. G., Kahn, T. G., Simcox, A., Schwartz, Y. B. & Pirrotta, V. Genome-wide activities of Polycomb complexes control pervasive transcription. Genome Res 25, 1170–1181, doi:10.1101/gr.188920.114 (2015).

77 Moussa, H. F. et al. Canonical PRC1 controls sequence-independent propagation of Polycomb-mediated gene silencing. Nat Commun 10, 1931, doi:10.1038/s41467-019-09628-6 (2019).

78 Blackledge, N. P. et al. PRC1 Catalytic Activity Is Central to Polycomb System Function. Mol Cell, doi:10.1016/j.molcel.2019.12.001 (2019).

79 Tamburri, S. et al. Histone H2AK119 Mono-Ubiquitination Is Essential for Polycomb-Mediated Transcriptional Repression. Mol Cell, doi:10.1016/j.molcel.2019.11.021 (2019).

80 Wu, R. S., Kohn, K. W. & Bonner, W. M. Metabolism of ubiquitinated histones. J Biol Chem 256, 5916–5920 (1981).

81 Joo, H. Y. et al. Regulation of cell cycle progression and gene expression by H2A deubiquitination. Nature 449, 1068–1072, doi:10.1038/nature06256 (2007).

82 Grau, D. J. et al. Compaction of chromatin by diverse Polycomb group proteins requires localized regions of high charge. Genes Dev 25, 2210–2221, doi:10.1101/gad.17288211 (2011).

83 Lau, M. S. et al. Mutation of a nucleosome compaction region disrupts Polycomb-mediated axial patterning. Science 355, 1081–1084, doi:10.1126/science.aah5403 (2017).

84 Beh, L. Y., Colwell, L. J. & Francis, N. J. A core subunit of Polycomb repressive complex 1 is broadly conserved in function but not primary sequence. Proc Natl Acad Sci U S A 109, E1063–1071, doi:10.1073/pnas.1118678109 (2012).

85 Berndsen, C. E. & Wolberger, C. A spectrophotometric assay for conjugation of ubiquitin and ubiquitin-like proteins. Anal Biochem 418, 102–110, doi:10.1016/j.ab.2011.06.034 (2011).

86 Brzovic, P. S., Lissounov, A., Christensen, D. E., Hoyt, D. W. & Klevit, R. E. A UbcH5/ubiquitin noncovalent complex is required for processive BRCA1-directed ubiquitination. Mol Cell 21, 873–880, doi:10.1016/j.molcel.2006.02.008 (2006).

87 Dyer, P. N. et al. Reconstitution of nucleosome core particles from recombinant histones and DNA. Methods Enzymol 375, 23–44, doi:10.1016/s0076-6879(03)75002-2 (2004).

88 Luger, K., Rechsteiner, T. J. & Richmond, T. J. Preparation of nucleosome core particle from recombinant histones. Methods Enzymol 304, 3–19, doi:10.1016/s0076-6879(99)04003-3 (1999).

89 Abmayr, S. M., Yao, T., Parmely, T. & Workman, J. L. Preparation of nuclear and cytoplasmic extracts from mammalian cells. Curr Protoc Pharmacol Chapter 12, Unit12 13, doi:10.1002/0471141755.ph1203s35 (2006).

90 Lee, K. M., Sif, S., Kingston, R. E. & Hayes, J. J. hSWI/SNF disrupts interactions between the H2A N-terminal tail and nucleosomal DNA. Biochemistry 38, 8423–8429, doi:10.1021/bi990090o (1999).

91 Carruthers, L. M., Tse, C., Walker, K. P., 3rd & Hansen, J. C. Assembly of defined nucleosomal and chromatin arrays from pure components. Methods Enzymol 304, 19–35, doi:10.1016/s0076-6879(99)04004-5 (1999).

92 Wong, I. & Lohman, T. M. A double-filter method for nitrocellulose-filter binding: application to protein-nucleic acid interactions. Proc Natl Acad Sci U S A 90, 5428–5432, doi:10.1073/pnas.90.12.5428 (1993).

93 Kachaner, D. et al. Coupling of Polo kinase activation to nuclear localization by a bifunctional NLS is required during mitotic entry. Nat Commun 8, 1701, doi:10.1038/s41467-017-01876-8 (2017).

94 Peng, K., Radivojac, P., Vucetic, S., Dunker, A. K. & Obradovic, Z. Length-dependent prediction of protein intrinsic disorder. BMC Bioinformatics 7, 208, doi:10.1186/1471-2105-7-208 (2006).

